# The chirality of the mitotic spindle provides a mechanical response to forces and depends on microtubule motors and augmin

**DOI:** 10.1101/2020.12.27.424486

**Authors:** Monika Trupinić, Barbara Kokanović, Ivana Ponjavić, Ivan Barišić, Siniša Šegvić, Arian Ivec, Iva M. Tolić

## Abstract

Forces produced by motor proteins and microtubule dynamics within the mitotic spindle are crucial for proper chromosome segregation. In addition to linear forces, rotational forces or torques are present in the spindle, reflected in the left-handed twisted shapes of microtubule bundles that make the spindle chiral. However, the biological role and molecular origins of spindle chirality are unknown. By developing methods for measuring spindle twist, we show that spindles are most chiral near the metaphase-to-anaphase transition. To assess the role of chirality in maintaining spindle robustness under force, we compressed the spindles along their axis. This resulted in stronger left-handed twist, suggesting that the twisted shape allows for a mechanical response to forces. Inhibition or depletion of motor proteins that perform chiral stepping, Eg5/kinesin-5, Kif18A/kinesin-8, MKLP1/kinesin-6, and dynein, decreased the left-handed twist or led to right-handed twist, implying that these motors regulate the twist by rotating microtubules within their antiparallel overlaps or at the spindle pole. Right-handed twist was also observed after the depletion of the microtubule nucleator augmin, indicating its contribution to the twist through the nucleation of antiparallel bridging microtubules. The uncovered switch from left-handed to right-handed twist reveals the existence of competing mechanisms that promote twisting in opposite directions. As round spindles were more twisted than elongated ones, we infer that bending and twisting moments are generated by similar molecular mechanisms and propose a physiological role for spindle chirality in allowing the spindle to absorb mechanical load.

## INTRODUCTION

Mitosis is a stage of the cell cycle in which replicated chromosomes are separated into two new nuclei destined to the two daughter cells^1^. In order to segregate the genetic material, the cell forms a mitotic spindle, a complex micro-structure made of microtubules and numerous associated proteins^2–4^. The spindle physically separates the chromosomes to the opposite poles of the cell and ensures that each daughter cell has the same number of chromosomes as the parental cell.

The spindle is a mechanical structure that can generate and balance forces within itself^5^. Forces in the spindle are crucial for proper spindle functioning in each phase of mitosis. Kinetochore fibers exert forces necessary for the positioning of the chromosomes at the center of the spindle in metaphase^6–8^ and for pulling the chromosomes apart during anaphase^9–11^. On the other hand, overlap bundles balance the forces at kinetochores by acting as bridges between sister kinetochore fibers in metaphase and anaphase^12–15^, and also regulate pole separation in anaphase^16–18^. All the forces in the spindle arise from active processes of motor proteins as well as microtubule polymerization and depolymerization^19–21^. Direct measurement of the forces in the spindle, although possible^22^, is challenging because of the small scales involved.

Forces are also responsible for the shape of a spindle. Due to the mechanical properties of microtubules, which are thin and elastic filaments that are inherently straight and curve under forces, the spindle obtains its characteristic shape^12,23,24^. This means that the spindle shape reflects the forces within it, which allows for an indirect measurement of forces by inferring them from the shapes of the microtubule bundles^12,25^, similarly to studies of forces and shapes of individual microtubules *in vitro* ^26,27^.

Recently, it was shown that the shape of the mitotic spindle in human HeLa and U2OS cells is chiral, as the spindle has a left-handed twist around the pole-to-pole axis^28^. Microtubule bundles twist because of the torques that exist within them in addition to linear forces. The experimentally measured three-dimensional shapes of the microtubule bundles, primarily bridging fibers which laterally link sister kinetochore fibers and are marked by protein regulator of cytokinesis 1 (PRC1), were used to deduce forces and torques in the spindle by comparison with a theoretical model^28^. Left-handed twist was also observed in spindles lacking NuMA and kinesin-5 activity in RPE1 cells during anaphase^29^. Another organism whose spindles are prominently twisted is a unicellular eukaryote, amoeba *Naegleria gruberi*. Interestingly, amoeba’s spindles are predominantly twisted in a right-handed fashion^30^.

The twist of the spindle is potentially generated by motor proteins that, in addition to linear forces, exert rotational forces on microtubules by switching protofilaments with a bias in a certain direction^31–42^. The first molecular motor discovered to generate torque was the single-headed axonemal dynein. In *in vitro* gliding motility assays, surface-attached dynein motors rotated the microtubules around their axis in a clockwise motion, when viewed from the minus-end of the microtubules, while translocating them in a linear fashion^31^. The same type of assay was used to show that the minus-end directed motor kinesin-14 (Ncd) generates torques which rotate microtubules in a clockwise direction as viewed from their minus-ends^32^, and similar results were obtained with kinesin-14 in an assay in which microtubules glide along each other, where the transport microtubule moves in a helical motion in a clockwise direction^41^.

Counterclockwise rotation direction has been found for the plus-end-directed motor protein kinesin-5 (Eg5), which was observed as a corkscrew motion of a sliding microtubule on surface-attached motors^34^. Gliding motility assays, as well as motility assays on freely suspended microtubules, showed a counterclockwise rotation for the plus-end-directed motor protein kinesin-8 (Kip3)^36,40^, while another study found that kinesin-8 can switch protofilaments in both directions^38^. Several other motor proteins also exhibit rotational movements, including kinesin-1^33,39^, kinesin-2^35^, cytoplasmic dynein^37^, and kinesin-6 (MKLP1)^42^. However, in contrast to this large body of knowledge on chiral motor stepping *in vitro*, the role of motor proteins and their asymmetric stepping in the generation of torques within microtubule bundles *in vivo* and consequently spindle twist is unknown.

In this paper we address the biological role and the molecular origin of spindle chirality. We show that spindle twist changes through different phases of mitosis and peaks around anaphase onset in a cancer and a non-cancer cell line. To test the idea that the chiral shape may help the spindle to absorb mechanical load, we compressed vertically oriented spindles along the pole-to-pole axis, which led to an increase in spindle twist. Thus, we propose a biological function of spindle chirality in promoting the flexibility of the spindle and its mechanical response to external forces, thereby decreasing the risk of spindle breakage under high load. By performing a candidate screen in which we depleted or inactivated motor proteins that step in a chiral manner and other microtubule-associated proteins, we identified several molecular players involved in the regulation of spindle chirality, leading us to suggest that the main mechanism generating spindle chirality is the action of motor proteins that rotate microtubules around one another within the antiparallel overlaps.

## RESULTS

### Spindle twist is most pronounced at anaphase onset in a cancer and a non-cancer cell line

To explore the twist of the spindle, the first step was to obtain end-on view images covering the whole spindle from pole to pole because this view allows for the visualization of the twist of microtubule bundles (Figure 1A, end-on view; Movie S1). A signature of the twisted shape is that microtubule bundles look like flower petals in the end-on view. In contrast, the twisted shape is not easily recognized in the side-view of the spindle (Figure 1A, side view). If the spindle is standing vertically with respect to the imaging plane, a z-stack of images directly provides an end-of view of the spindle, but if the spindle is lying horizontally, a z-stack needs to be transformed into the end-on view.

**Figure 1.**
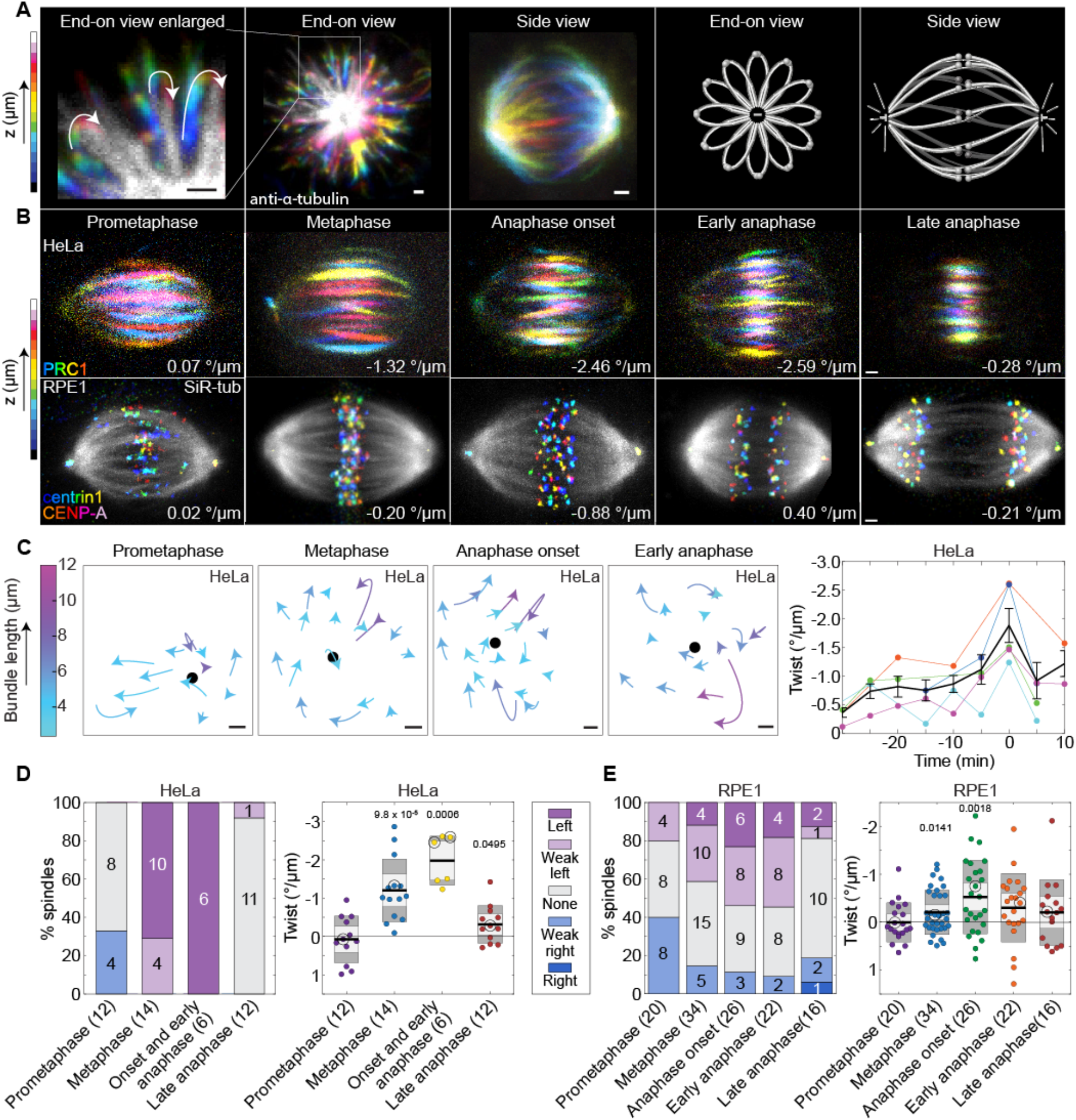
Spindle twist culminates at the beginning of the anaphase. **(A)** Images of spindles immunostained for α-tubulin in a HeLa-Kyoto BAC cell line stably expressing PRC1-GFP (PRC1-GFP signal is not shown). Left, enlarged section of the image showing microtubule bundles from the end-on view, rotating in a clockwise direction (arrows) through z-planes when moving towards the observer. The end-on view and the side view of a spindle are shown next. Images are color-coded for depth using ImageJ temporal color coding Lookup Table ‘16 colors’ (see color bar). Scale bar, 1 μm. Right two panels show schemes of a spindle from the end-on view and side view. **(B)** Top row, images of spindles in HeLa-Kyoto BAC cells stably expressing PRC1-GFP, shown in different phases of mitosis. Bottom row, images of spindles in hTERT-RPE1 cells expressing CENP-A-GFP and centrin1-GFP, shown in different phases of mitosis. Microtubule bundles of HeLa spindles (PRC1-GFP signal) are color-coded for depth (every second plane was used) using ImageJ temporal color coding Lookup Table ‘16 colors’ and microtubules of RPE1 spindles are shown in grey (SiR-tubulin dye) while kinetochores/centrosomes are color-coded for depth using the same color coding as described above (see color bar) and filtered with a Gaussian blur (radius 0.7). Scale bars, 1 μm. Additional examples of HeLa cell spindles are shown in Figures S1C and S1D-F. **(C)** On the left, representation of microtubule bundle movements along the z-axis of the same spindle in HeLa cells viewed from the end-on view in different phases of mitosis; each microtubule bundle is represented by a circular arc of the circle fitted on the traces with arrowheads pointing in the rotation direction, the color of the arrow shows the bundle length along the pole-to-pole axis (shared color bar on the left); black dot represents the pole-to-pole axis; scale bar, 1 μm. On the right, graph shows the change of twist values for five HeLa cells over time; the beginning of anaphase (visible chromosome separation) was set as time zero; each color represents one cell and thick black line represents mean values with error bars showing SEM; data shown in orange originate from the cell whose bundles’ movements are shown on the left and images are shown in Fig. S1C. Experiments were performed on the HeLa-Kyoto BAC cells stably expressing PRC1-GFP. Other examples of spindle images are shown in Figure S1C. **(D)** Twist in different phases of mitosis in HeLa-Kyoto BAC cells stably expressing PRC1-GFP. On the left, visual assessment graph represents percentages of spindles showing left, right, weak left, weak right or no twist, as described in the legend, in different phases of mitosis (numbers on the graph and in the brackets show the number of cells). On the right, graph shows twist values calculated with the optical flow method in different phases of mitosis. The black line shows the mean; the light and dark grey areas mark 95% confidence interval on the mean and standard deviation, respectively; numbers above the data show p-values (Student’s t-test for the mean twist value different from 0). Non-significant differences are not shown. Circled dots represent the cells that are shown on the images above. Raw data of 10 out of 14 metaphase spindles was taken and re-calculated from^28^ and also used in Figure S1B. **(E)** Twist in different phases of mitosis in hTERT-RPE1 cells expressing CENP-A-GFP and centrin1-GFP; legend as in (D).

To quantify spindle twist, we used 3 complementary approaches: visual assessment, optical flow, and bundle tracing (Figure S1A; Methods). As it is still an open question in the field what method is the most appropriate to measure spindle twist^28,29,43^, visual assessment is useful as a quick and rough estimate of the twist and as a control for automated or semi-automated methods. In the visual assessment method, the spindle is observed end-on and the rotation of microtubule bundles around the pole-to-pole axis is estimated visually. If the bundles rotate clockwise when moving along the spindle axis in the direction towards the observer, the twist is left-handed, and vice versa (Figure S1A, left). We score the twist as left-handed, right-handed, weak left-handed, weak right-handed, or no visible twist. Weak twists correspond to a range of approximately −1 to −2 °/μm in the bundle tracing method (Figure S1B). This is visible as a total rotation of 5-10° in the clockwise (left-handed) or counter-clockwise (right-handed) direction in the end-on view of the spindle when moving towards the observer along the bundle length, which is typically 5 μm. Accordingly, left and right twists correspond to a rotation of more than 10° in the end-on view (Methods; Figure S1B). In the optical flow method, the movement of the signal coming from microtubule bundles is estimated by comparing the signal from one z-plane to the next (Figure S1A, middle; Methods). This method provides a value for the average twist of all bundles in a spindle and is optimal for experiments on a large number of spindles because it is automated. The bundle tracing method is an extension of the approach developed previously^28^, where individual bundles are manually traced by following the bundle contour in the end-on view of the spindle (Figure S1A, right; Methods). Subsequently, each bundle is analyzed so that a plane is fitted to the points representing the bundle, and a circle that lies within this plane is fitted to these points and used to calculate the curvature and twist of individual bundles^43^ (Figure S1A, right; Methods).

As a label for microtubule bundles, we used SiR-tubulin to observe all microtubule bundles, or PRC1-GFP to observe the bridging fibers^12,13^. To compare the results of the three methods, we analyzed twist of 10 metaphase spindles in HeLa cells stably expressing PRC1-GFP (Figure S1B). All three methods yielded a left-handed twist, which is expressed by negative values, for all 10 spindles. The spindles that were visually assessed as having a strong left twist had, on average, a higher left twist value also in the bundle tracing and optical flow method, than those with a weak left twist. The absolute values of twist of individual spindles obtained by bundle tracing and optical flow were similar, with optical flow yielding smaller negative values (−1.32 ± 0.29 °/μm, n = 10; all data are given as mean ± sem) than bundle tracing (−2.07 ± 0.29 °/μm, n = 10). This difference is likely due to the sensitivity of the optical flow method to all signal including the background. Based on this cross-check between the three methods, we conclude that they provide comparable values of spindle twist. Thus we use optical flow for experiments in which we test changes in the overall spindle twist in a large number of cells, and bundle tracing for experiments where high spatial precision is required.

Spindles in cancer cell lines are twisted in a left-handed manner in metaphase^28^, but it is not known whether the twist is present already when the spindle assembles in prometaphase or it arises as the spindle matures. Furthermore, it is unknown how the twist changes during anaphase. To examine the development of spindle twist throughout mitosis (Figure 1B; Movies S2-S6), we first measured twist in individual live HeLa cells expressing PRC1-GFP as they progressed through mitosis (Figure 1C; Figure S1C). The average twist of the spindle in prometaphase was close to 0, it was left-handed (negative) during metaphase, culminated at anaphase onset reaching a value of −1.88 ± 0.3 °/μm (n = 5), and decreased afterwards (Figure 1C). In agreement with this result, experiments in which different spindles were imaged in different phases showed a peak of spindle twist around anaphase onset, with a value of −1.98 ± 0.26 °/μm (n = 6) (Figures 1B, 1D; Figures S1D-F; Movies S2-S6; Table 1). Expression of PRC1-GFP in this cell line did not influence the twist, as non-transfected HeLa cells stained with SiR-tubulin showed similar twist values in metaphase (Table 1; p = 0.47, Student’s t-test).

**Table 1.** Spindle twist, length and width in HeLa and RPE1 cells in different phases of mitosis and after protein perturbations. All values are shown as mean ± s.e.m. Purple color denotes left-handed twist (mean value of twist < 0 and p < 0.05 for a difference from 0 in a t-test), blue color denotes right-handed twist (mean value of twist > 0 and p < 0.05 for a difference from 0 in a t-test), and grey denotes no twist (p > 0.05 for a difference from 0 in a t-test). The numbers in the brackets denote the number of cells; n.d., not determined; * represents nontransfected HeLa cells (the rest of the data on HeLa cells comes from HeLa-Kyoto BAC cells stably expressing PRC1-GFP); RPE1 cells used were hTERT-RPE1 cells permanently transfected and stabilized using CENP-A-GFP and centrin1-GFP.

|  | HeLa |  |  | RPE1 |  |  |
| --- | --- | --- | --- | --- | --- | --- |
| | Twist ($^{\circ}/\mu\text{m}$) | Length ($\mu\text{m}$) | Width ($\mu\text{m}$) | Twist ($^{\circ}/\mu\text{m}$) | Length ($\mu\text{m}$) | Width ($\mu\text{m}$) |
| Prometaphase | $0.09 \pm 0.18$<br>(12) | $11.8 \pm 0.2$ | $8.3 \pm 0.3$ | $0.004 \pm 0.09$ (20) | $12.1 \pm 0.2$ | $8.7 \pm 0.1$ |
| Metaphase | $-1.20 \pm 0.22$ (14) | $11.5 \pm 0.3$ | $9.0 \pm 0.2$ | $-0.21 \pm 0.08$ (34) | $12.8 \pm 0.3$ | $9.0 \pm 0.1$ |
| | $*-1.01 \pm 0.14$ (14) | $*10.6 \pm 0.1$ | $*9.9 \pm 0.2$ | | | |
| Anaphase onset | $-1.98 \pm 0.26$ (6) | $12.2 \pm 0.4$ | $9.0 \pm 0.5$ | $-0.53 \pm 0.15$ (26) | $12.9 \pm 0.2$ | $8.7 \pm 0.1$ |
| Early anaphase | | | | $-0.30 \pm 0.15$ (22) | $13.8 \pm 0.3$ | $8.4 \pm 0.2$ |
| Late anaphase | $-0.31 \pm 0.14$ (12) | $13.3 \pm 0.2$ | $9.0 \pm 0.3$ | $-0.20 \pm 0.17$ (16) | $16.6 \pm 0.4$ | $7.5 \pm 0.3$ |
| Mps1 inhibition | $-0.17 \pm 0.21$ (17) | $11.9 \pm 0.4$ | $8.4 \pm 0.2$ | n.d. | n.d. | n.d. |
| Eg5 inhibition (after $< 5$ min) | $-0.47 \pm 0.14$ (16) | $12.0 \pm 0.2$ | $9.3 \pm 0.2$ | $-0.06 \pm 0.19$ (11) | $12.3 \pm 0.4$ | $8.8 \pm 0.1$ |
| Eg5 inhibition (after 10-20 min) | n.d. | n.d. | n.d. | $0.06 \pm 0.13$ (12) | $8.3 \pm 0.2$ | $7.7 \pm 0.2$ |
| Eg5 overexpression | n.d. | n.d. | n.d. | $-0.25 \pm 0.12$ (11) | $12.7 \pm 0.4$ | $9.0 \pm 0.3$ |
| Kif18A siRNA | $0.11 \pm 0.14$ (21) | $12.8 \pm 0.6$ | $8.3 \pm 0.2$ | $0.30 \pm 0.11$ (24) | $15.0 \pm 0.6$ | $8.7 \pm 0.2$ |
| Kif18A overexpression | n.d. | n.d. | n.d. | $-0.26 \pm 0.20$ (7) | $10.3 \pm 0.3$ | $8.1 \pm 0.2$ |
| MKLP1 siRNA | $-0.80 \pm 0.19$ (12) | $12.1 \pm 0.1$ | $8.4 \pm 0.2$ | $0.48 \pm 0.10$ (16) | $13.6 \pm 0.3$ | $9.1 \pm 0.3$ |
| HSET siRNA | $-1.13 \pm 0.21$ (17) | $11.7 \pm 0.2$ | $8.4 \pm 0.2$ | $-0.19 \pm 0.12$ (18) | $13.9 \pm 0.4$ | $8.9 \pm 0.1$ |
| Dynein inhibition | $-0.18 \pm 0.11$ (18) | $9.8 \pm 0.1$ | $8.3 \pm 0.2$ | $-0.18 \pm 0.08$ (16) | $9.8 \pm 0.3$ | $8.4 \pm 0.1$ |
| Dynein KO | n.d. | n.d. | n.d. | $-0.29 \pm 0.13$ (15) | $15.8 \pm 0.8$ | $10.7 \pm 0.5$ |
| PRC1 siRNA | *-0.94 ± 0.17 (19) | *9.9 ± 0.2 | *9.9 ± 0.1 | 0.22 ± 0.11 (22) | 15.3 ± 0.4 | 9.6 ± 0.2 |
| PRC1 overexpression | n.d. | n.d. | n.d. | -0.08 ± 0.11 (10) | 10.3 ± 0.4 | 8.0 ± 0.1 |
| HAUS6 siRNA | 0.18 ± 0.21 (16) | 11.9 ± 0.3 | 9.4 ± 0.4 | 0.49 ± 0.21 (16) | 11.7 ± 0.3 | 9.0 ± 0.1 |
| HAUS8 siRNA | -0.35 ± 0.40 (10) | 12.1 ± 0.4 | 9.6 ± 0.5 | 0.85 ± 0.24 (13) | 13.1 ± 0.4 | 9.0 ± 0.2 |
| Mock siRNA | *-0.85 ± 0.20 (17) | *10.7 ± 0.3 | *9.6 ± 0.2 | -0.22 ± 0.08 (39) | 12.5 ± 0.2 | 8.6 ± 0.1 |
|  | -0.94 ± 0.16 (13) | 11.2 ± 0.3 | 9.5 ± 0.3 |  |  |  |
| MG-132 | n.d. | n.d. | n.d. | 0.51 ± 0.14 | 12.0 ± 0.4 | 8.8 ± 0.2 |

To test whether the time spent in metaphase affects spindle twist, we accelerated entry into anaphase by inhibiting Mps1 kinase, one of the main components of the spindle assembly checkpoint^44^. Treatment of HeLa cells expressing PRC1-GFP with the inhibitor AZ3146^45^ during prometaphase shortened the time to anaphase from 40 minutes to 8-10 minutes, on average. We measured the twist at the beginning of anaphase or in early anaphase and found it to be significantly smaller than in control cells, −0.17 ± 0.21 °/μm (n = 17) (Figure S1G; Table 1). This result suggests that the reduction of time needed to enter the anaphase may also mean a reduction of time to build up spindle twist.

To explore whether spindle twist and its variation over time is specific to cancer cell lines, we measured twist in the non-cancer immortalized epithelial cell line hTERT-RPE1 (from here on referred to as RPE1) (Figure 1B) and found that spindles in these cells also show a left-handed twist, but the values are smaller than in HeLa cells (Figure 1E). Moreover, the temporal pattern of twist in RPE1 cells was similar to that in HeLa cells. Twist was absent in prometaphase, it was very weak left-handed in metaphase, had a peak value at anaphase onset, decreased during anaphase, and vanished in late anaphase (Figure 1E; Movies S2-S6; Table 1). The value at anaphase onset was −0.53 ± 0.15 °/μm (n = 26), which indicates a weaker left-handed twist than in HeLa cells. Taken together, our results show that spindles are born without twist. The left-handed twist in HeLa cells arises as the spindle acquires its metaphase shape, peaks at the start of chromosome segregation, and declines afterwards. In RPE1 cells, the twist shows a similar trend, but the values are much less pronounced and the twist is mostly noticeable only in early anaphase.

### Compression of the spindle along the pole-to-pole axis increases the left-handed twist

The biological role of spindle chirality is still unknown. Although chirality may be simply a side effect of the activity of torque-generating motors, the twisted shapes of microtubule bundles may contribute to spindle physiology by allowing changes of spindle shape as a mechanical response to external forces. To test this idea, we gently compressed vertically oriented HeLa cell spindles in metaphase along the pole-to-pole axis for 1.5 minute, following the compression protocol from a previous study^46^ (Figure 2A; Movie S7). We used the bundle tracing method to measure spindle twist, which allowed us to graphically reconstruct spindles from the end-on view and side view (Figure 2B). Traces of the microtubule bundles in the end-on view after 1 minute of compression were more rounded than before compression, indicating an increase in twist, and the mitotic spindle shortened (Figure 2B). Spindle shortening was used as a measure to confirm successful compression, showing that spindle length decreased from 14.07 ± 0.55 μm before compression to 12.75 ± 0.80 μm after 1 minute of compression (p = 0.013; a paired t-test was used to compare the values before and after compression, n = 6 spindles) (Figure 2C). Spindle width increased after compression in some cases, e.g., for the spindle shown in Figure 2B, but overall this change was not significant (p = 0.18) (Figure 2D).

**Figure 2.**
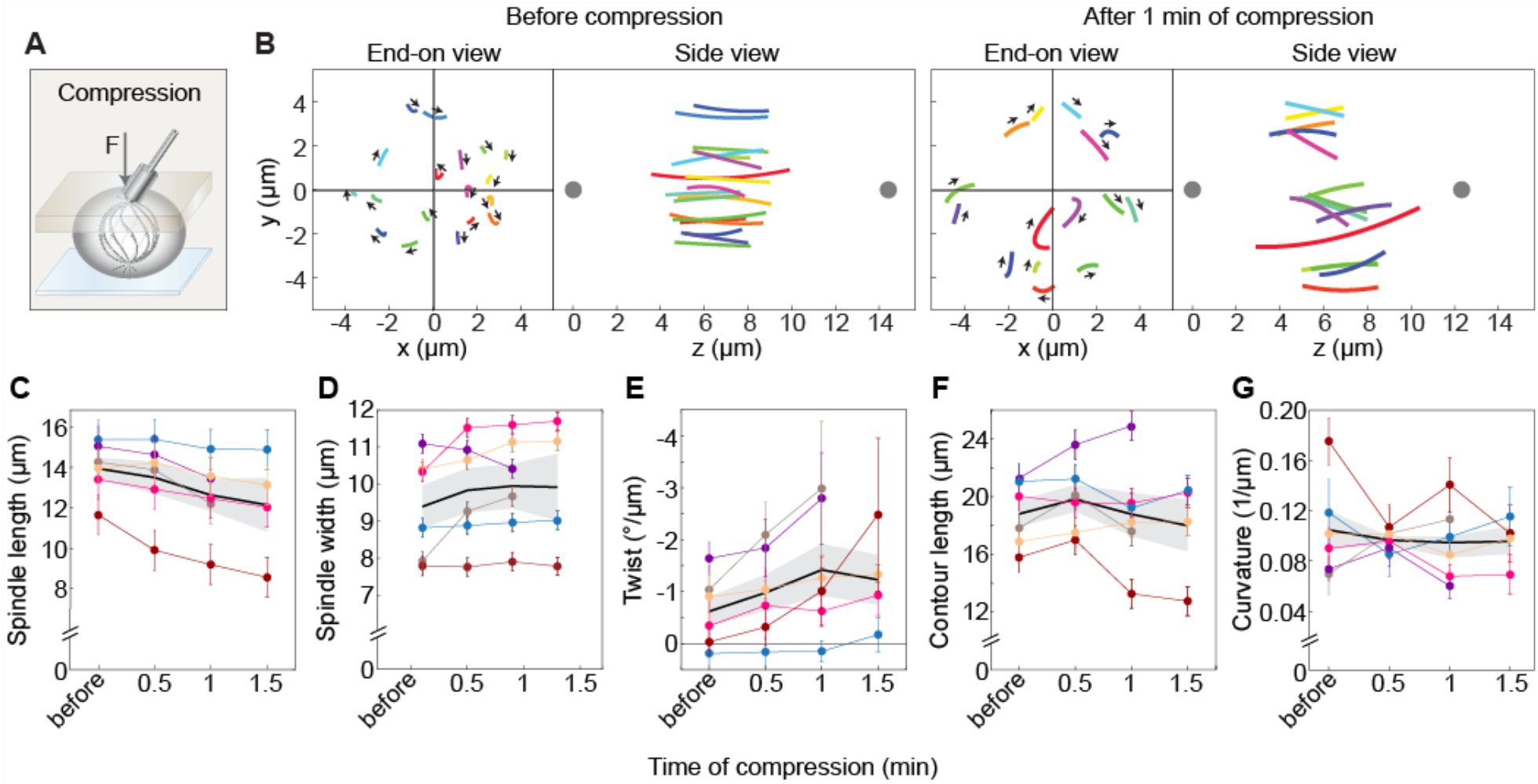
Spindles compressed by an external force have stronger twist. **(A)** Scheme of the experimental method for spindle compression. Blue layer represents the dish; spindle is shown inside a cell with microtubule bundles in grey; grey layer represents the gel with a metal rod on top; arrow shows the direction of force. (**B**) Microtubule bundles in a spindle shown from the end-on view and side view before compression and after 1 minute of compression, as indicated. Each bundle is represented by a different color (in the end-on view and side view, individual bundles are colored with the same color, but colors before compression and after 1 min of compression do not represent the same bundles); lines show circular arcs of the fitted circles and arrows represent the rotation direction of each bundle. Grey dots in the side view represent spindle poles. **(C)-(G)** Graphs showing the change of spindle parameters from before compression up to 1.5 min of compression: spindle length **(C)**, spindle width (**D**), twist of microtubule bundles **(E)**, length of the bundle contours **(F)**, and bundle curvature **(G)**. Each color represents one cell; dots represent mean values; error bars in (C) and (D) show the estimated errors in the determination of spindle length and width, 1 μm and 0.25 μm, respectively; error bars in other graphs represent SEM. Thick black line shows the mean of all data with grey area representing SEM. Experiments were performed on the HeLa-Kyoto BAC cells stably expressing PRC1-GFP. Successful compression was performed on 6 spindles from 5 independent experiments. Individual data points are shown in Figure S2A.

Interestingly, compression resulted in a 2.3-fold increase of the left-handed spindle twist, from −0.63 ± 0.28 °/μm before compression to −1.42 ± 0.50 °/μm after 1 minute of compression (p = 0.040) (Figure 2E; Figure S2A; Movie S7). Histograms of twist values show that the distribution shifted towards more negative values upon compression (Figure S2B). To quantify this shift, we analyzed the fraction of bundles having a strong left-handed twist with a value smaller than −2.8 °/μm, which is one standard deviation away from the mean twist before compression. The twist was smaller than −2.8 °/μm for 9 out of 80 bundles (11.3% ± 3.5%) before compression, whereas after compression this was the case for 21 out of 73 bundles (28.8% ± 5.3%). The difference was significant (p = 0.0064; two-proportions Z-test), suggesting that compression resulted in a higher proportion of bundles having a strong left-handed twist.

Contour length of the microtubule bundles did not change significantly after compression (p = 0.99) (Figure 2F; Figure S2A). We were unable to detect changes in bundle curvature after compression (p = 0.41), (Figure 2G; Figure S2A), which is consistent with the non-significant change in spindle width. Thus, as the spindle was compressed end-on by an external force, which resulted in spindle shortening, the microtubule bundles did not shorten substantially, but instead became more twisted. These results support the idea that the twist within the bundles allows a mechanical response to external forces.

### Motor proteins Eg5/kinesin-5, Kif18A/kinesin-8, MKLP1/kinesin-6, and dynein regulate spindle twist

To explore the molecular origins of torques in the spindle and thus its twisted shape, we consider the following molecular activities. First, motors that exert torque on the microtubule may generate the twisted shape of the bundle by twisting the microtubules within the bundle around each other, or by twisting the microtubules with respect to the spindle pole. Second, proteins that crosslink neighboring microtubules or link microtubules with the pole may prevent free rotation of the microtubules, thereby allowing for twisting of the bundles. Third, nucleation of new microtubules within the bundle may affect the bundle twist.

To test the role of these activities in the regulation of spindle twist, we performed a candidate screen on HeLa and RPE1 cells in which we perturbed motor proteins and other microtubule-associated proteins one by one using siRNA-mediated depletion, small-molecule inhibitors, or overexpression, and measured the resulting spindle twist. As the candidates for this mini-screen, we selected spindle-localized motor proteins for which it has been shown *in vitro* that they can rotate the microtubule (Eg5/kinesin-5, Kif18A/ kinesin-8, MKLP1/kinesin-6, HSET/kinesin-14, dynein), the main crosslinker of antiparallel microtubules PRC1, and the augmin complex that is responsible for the nucleation of microtubules along existing microtubules. Spindle twist was measured during metaphase, rather than at anaphase onset when twist is most pronounced, because depletion or inhibition of some of the candidate proteins, such as Eg5, Kif18A, and augmin, interferes with anaphase entry^47,48 49^. Furthermore, the measurement of the twist in metaphase is more reproducible because spindles in metaphase are in a steady state, whereas anaphase spindles undergo extensive changes. All candidate proteins were depleted by siRNA, except Eg5 and dynein. Eg5 was inhibited with S-trityl-L-cysteine (STLC)^50^, because siRNA depletion of Eg5 would not allow for spindles to properly assemble, resulting in monoasters^49^. For dynein inhibition we used dynarrestin^51^ in both HeLa and RPE1 cells, as well as CRISPR/Cas9-inducible DYNC1H1 (dynein heavy chain) knockout (KO) RPE1 cells^52^. Depletion by siRNA of each protein was confirmed by measurements of the immunofluorescence signal of that protein on the spindle (Figures S3A, S3B).

In agreement with our previous work on HeLa cells^28^, we found that acute inhibition of Eg5 with STLC decreased the left-handed spindle twist in both HeLa and RPE1 cells (Figures 3A, 3B, 3C; Figures S4, S5; Table 1). In RPE1 cells, the spindles had no twist 5 minutes after STLC addition, while the spindle length was the same as before the treatment, and after 10-20 minutes, when the spindles were shorter but still bipolar (Table 1, Figure S4). These results suggest that changes in the spindle twist due to Eg5 inhibition are independent of the changes in spindle length.

**Figure 3.**
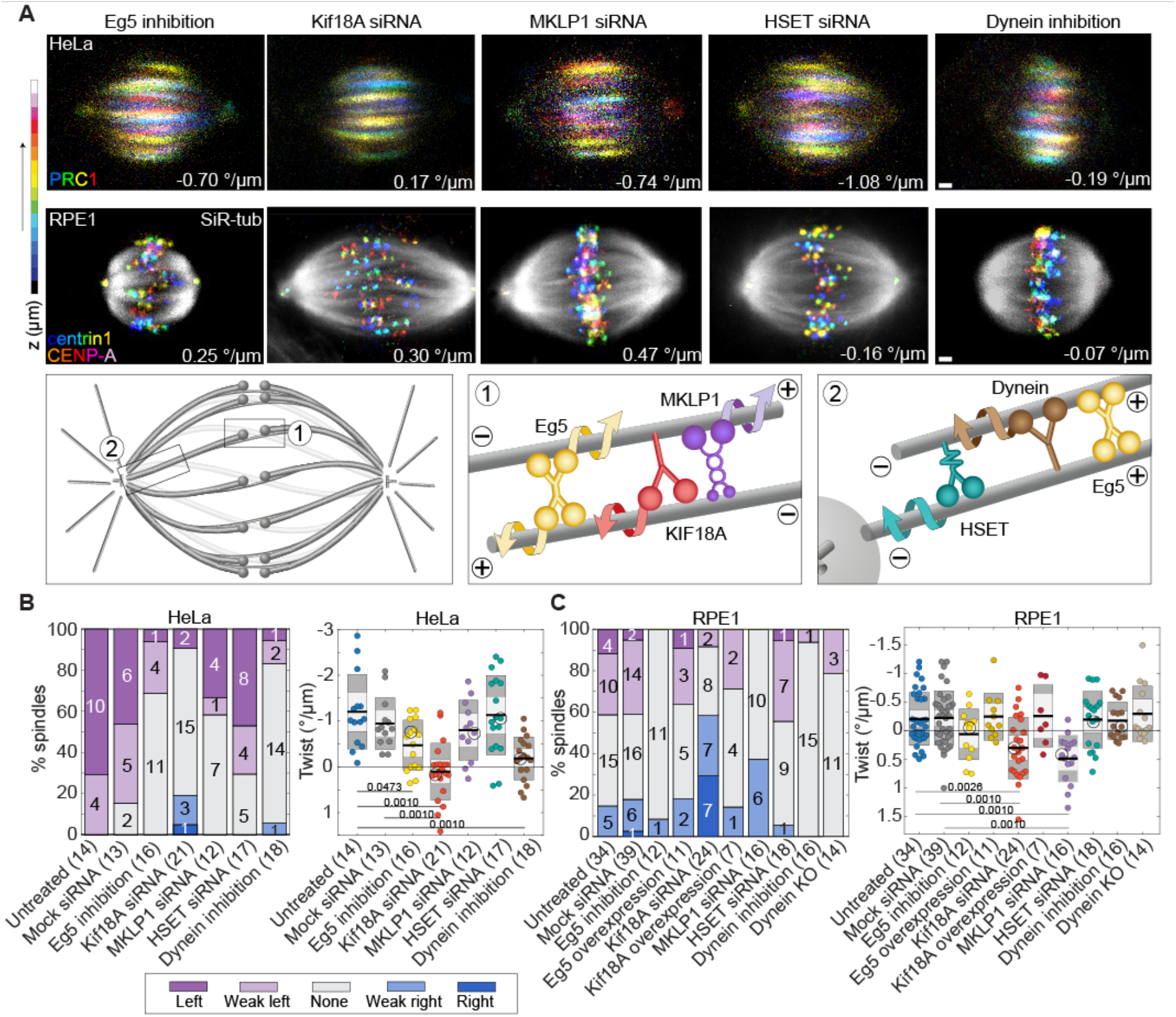
Motor proteins Eg5 and Kif18A control spindle twist. **(A)** First row, images of the spindles in HeLa-Kyoto BAC cells stably expressing PRC1-GFP after inhibition/depletion of Eg5, Kif18A, MKLP1, HSET, and dynein. Second row, images of the spindles in hTERT-RPE1 cells permanently transfected and stabilized using CENP-A-GFP and centrin1-GFP after inhibition/depletion of Eg5, Kif18A and HSET motor proteins. Third row, simplified schemes showing localization and movement of Eg5, Kif18A, MKLP1, HSET, and dynein in the spindle. Microtubule bundles of HeLa spindles (PRC1-GFP signal) are color-coded for depth (every second plane was used) by using ImageJ temporal color coding Lookup Table ‘16 colors’ and microtubules of RPE1 spindles are shown in grey (SiR-tubulin dye) while kinetochores/centrosomes are color-coded for depth using the same color coding as described above (see color bar) and filtered with a Gaussian blur (radius 0.7). Scale bars, 1 μm. Additional examples of spindles are shown in Figures. S4, S5. **(B)** Spindle twist after perturbations of motor proteins in HeLa cells expressing PRC1-GFP. Left, visual assessment graph represents percentages of spindles showing left, right, weak left, weak right or no twist, as described in the legend, after inhibition/depletion of Eg5, Kif18A, MKLP1, HSET, and dynein (numbers on the graph and in the brackets show the number of cells). Right, graph shows twist values calculated with the optical flow method after inhibition/depletion of Eg5, Kif18A, MKLP1, HSET, and dynein. The black line shows the mean; the light and dark grey areas mark 95% confidence interval on the mean and standard deviation, respectively; one-way ANOVA test showed significant difference between group means (p = 1.39 x 10^-8^); numbers below the data show p-values (Tukey’s HSD post hoc test); non-significant differences are not shown. Encircled dots represent cells on the images above. **(C)** Spindle twist after perturbations of motor proteins in RPE1 cells expressing CENP-A-GFP and centrin1-GFP, and RPE1 inducible CRISPR/Cas9 DYNC1H1 knockout cells; legend as in (B). Data for Eg5 inhibition correspond to 10-20 minutes after STLC addition. One-way ANOVA test showed significant difference between group means (p = 2.27 x 10^-7^).

Depletion of Kif18A abolished spindle twist in HeLa cells and resulted in right-handed twist in RPE1 cells, causing ~71% of RPE1 spindles to twist in the right-handed fashion, with a mean twist of 0.30 ± 0.11 °/μm (n = 24, p = 0.0119 for a difference from 0 in a Student’s t-test; Figures 3A, 3B, 3C; Figures S4, S5; Movie S8; Table 1). Overexpression of either Eg5 or Kif18A in RPE1 cells did not yield any changes in twist (Figure 3C; Figure S5; Table 1).

Depletion of MKLP1 did not change twist in HeLa cells, but significantly changed the twist in RPE1 cells, where 94% of spindles were twisted in a right-handed direction (Figures 3A, 3B, 3C; Figures S4, S5; Movie S9; Table 1). The mean twist was 0.48 ± 0.10 °/μm (n = 16, p = 0.0003 for a difference from 0 in a Student’s t-test).

Depletion of HSET/kinesin-14 did not change twist in HeLa or RPE1 cells (Figures 3A, 3B, 3C; Figures S4, S5; Table 1). Dynarrestin treatment abolished the twist of HeLa and RPE1 spindles, with a clearly visible effect on HeLa spindles, whereas the changes in RPE1 cells were subtle (Figures 3A, 3B, 3C; Figures S4, S5; Movie S10; Table 1). In DYNC1H1 knockout RPE1 cells the twist was also absent, but it was challenging to measure the twist in these cells due to the unfocused spindle poles and altered spindle shape (Figure 3C; Figure S5; Movie S10; Table 1). We conclude that Eg5, Kif18A, MKLP1, and dynein regulate the torques within the spindle, which lead to the twisted shape of microtubule bundles, and that their contribution differs in different cell lines.

### Depletion or overexpression of PRC1 in RPE1 spindles results in no twist

PRC1 protein is a key regulator of cytokinesis^53^, but also the main crosslinking protein of antiparallel microtubules within bridging fibers^12,13^. Without PRC1, bridging fibers are thinner and spindles have less curved and more diamond-like shape^12,54^, which led us to hypothesize that the twist might also be affected. In HeLa cells, depletion of PRC1 did not yield changes in the spindle twist (Figures 4A, 4B; Figure S4; Movie S9; Table 1). When we depleted PRC1 in RPE1 cells, the spindles had no twist, on average (Figures 4A, 4C; Figure S5; Movie S11; Table 1). Overexpression of PRC1 in RPE1 cells also resulted in the abolishment of the spindle twist, as the microtubule bundles became almost straight (Figures 4A, 4C; Figure S5; Movie S11; Table 1). These data suggest that PRC1 regulates torques within the spindle in RPE1 cells, possibly by limiting free rotation of microtubules within antiparallel bundles and by modulating the torsional rigidity of the bundle.

**Figure 4.**
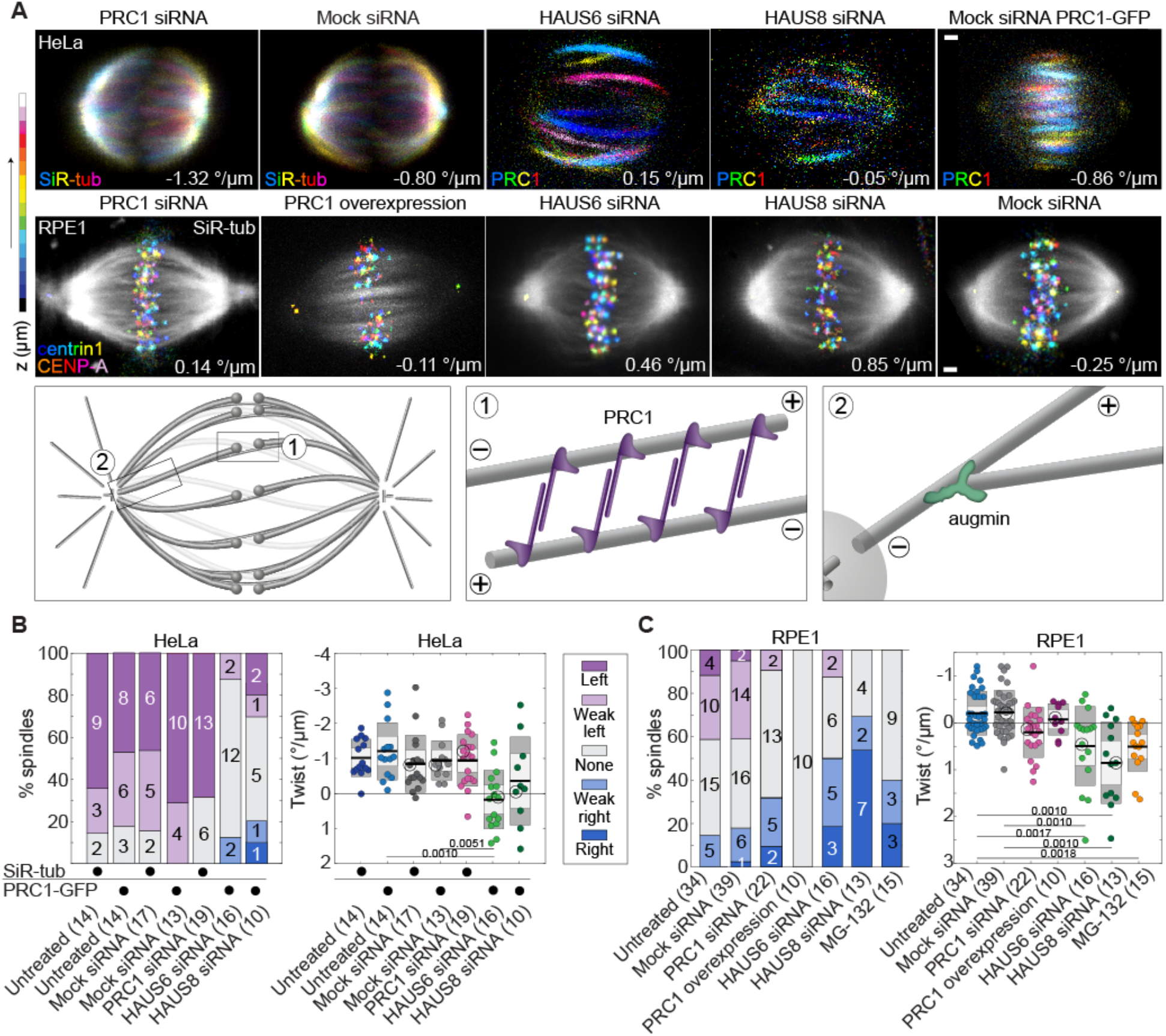
Microtubule crosslinker PRC1 and nucleator augmin regulate spindle twist. **(A)** First row, images of spindles in a non-transfected HeLa cell line stained with SiR-tubulin (first 2 spindles) and HeLa-Kyoto BAC cells stably expressing PRC1-GFP (last 3 spindles), after depletion of PRC1 and subunits of the augmin complex HAUS6 and HAUS8. Second row, images of spindles in the hTERT-RPE1 cells, permanently transfected and stabilized using CENP-A-GFP and centrin1-GFP RPE1, after perturbations of PRC1 and depletions of subunits of the augmin complex HAUS6 and HAUS8. Third row, simplified schemes showing the localization of PRC1 and augmin in the spindle. Microtubule bundles in HeLa cells (PRC1-GFP signal in HeLa-Kyoto BAC cells and SiR-tubulin dye in non-transfected HeLa cells) are color-coded for depth (every second plane was used) by using ImageJ temporal color coding Lookup Table ‘16 colors’ and microtubules of RPE1 spindles are shown in grey (SiR-tubulin dye, except the cells with overexpressed PRC1 that show PRC1-mCherry signal), while kinetochores/centrosomes are color-coded for depth using the same color coding as described above (see color bar) and filtered with a Gaussian blur (radius 0.7). Scale bars, 1 μm. Additional examples of spindles are shown in Figures S4, S5. **(B)** Spindle twist after perturbations of PRC1 and augmin in HeLa cells. Left, visual assessment graph represents percentages of spindles showing left, right, weak left, weak right or none twist, as described in the legend, after depletions of PRC1 protein and subunits of augmin complex HAUS6 and HAUS8 in HeLa cells (numbers on the graph and in the brackets show the number of cells; black dots under the graph denote the cell line/staining used for the corresponding treatment). Right, graph shows twist values calculated with the optical flow method after depletions of PRC1 protein and subunits of augmin complex HAUS6 and HAUS8. The black line shows the mean; the light and dark grey areas mark 95% confidence interval on the mean and standard deviation, respectively; one-way ANOVA test showed significant difference between group means (p = 8.06 x 10^-5^); numbers below the data show p-values (Tukey’s HSD post hoc test); nonsignificant differences are not shown. Encircled dots represent cells that are shown on images. Experiments were performed on the non-transfected HeLa cell line (for the depletion of PRC1 and its control) and HeLa-Kyoto BAC cells stably expressing PRC1-GFP (for the depletion of the HAUS6 and HAUS8 and their controls). **(C)** Spindle twist after perturbations of PRC1 and augmin in RPE1 cells expressing CENP-A-GFP and centrin1-GFP; legend as in (B). One-way ANOVA test showed significant difference between group means (p = 1.72 x 10^-9^).

### Depletion of augmin leads to no twist in HeLa cells and right-handed twist in RPE1 cells

The augmin complex is responsible for the microtubule nucleation from the lateral surface of the pre-existing microtubules^48,55^. Augmin is important for the nucleation of the bridging fibers and, consequentially, the maintenance of the spindle shape^56^. When we depleted the augmin subunit HAUS6 (hDgt6/FAM29A), which binds to γTuRC through the adaptor protein NEDD1^48^, the spindles in HeLa cells had zero twist on average, whereas those in RPE1 cells had right-handed twist of 0.49 ± 0.21 °/μm (n = 16, p = 0.0341 for a difference from 0 in a Student’s t-test; Figures 4A, 4B, 4C; Figures S4, S5; Table 1). A similar result was observed after the depletion of the augmin subunit HAUS8 (hDgt4/Hice1), which binds to pre-existing microtubules^57,58^. This resulted in zero average twist in HeLa cells and a strong right-handed twist in RPE1 cells of 0.85 ± 0.24 °/μm (n = 13, p = 0.0041 for a difference from 0 in a Student’s t-test; Figures 4A, 4B, 4C; Figures S4, S5; Movie S12; Table 1). The twist after depletion of HAUS6 or after depletion of HAUS8 was not significantly different in HeLa (p = 0.26) or RPE1 cells (p = 0.27), as expected given that they are part of the same complex. Thus, augmin-mediated nucleation of microtubules along the wall of pre-existing microtubules is an important determinant of the direction and amount of spindle twist.

As depletion of the augmin complex subunits prolongs metaphase^48^, we explored how the twist changes when cells are arrested in metaphase by adding the proteasome inhibitor MG-132. Interestingly, spindles in RPE1 cells arrested in metaphase had right-handed twist of 0.51 ± 0.14 °/μm (n = 15, Figure 4C; Figure S5; Table 1), suggesting that prolonging metaphase may cause a shift in the balance of torque-generating activities resulting in a right-handed twist.

### Round spindles are more twisted than elongated spindles

While imaging different types of spindles (phases of mitosis, cell lines, protein perturbation), we noticed that round spindles often have stronger twist than elongated spindles. To quantify this observation, we measured spindle length and width and calculated the width/length ratio as a measure for the roundness of the spindle, where ratios closer to 1 describe round spindles, and smaller values elongated spindles (Figure 5A, scheme). Higher width/length ratios are a signature of stronger bending moments in the spindle^28^. We tested the correlation between the width/length ratios and twist values in metaphase, and found that in non-transfected HeLa cells, whose width/length ratios were roughly between 0.8 and 1, rounder spindles had a stronger left-handed twist (Figure 5A), indicating a correlation between bending and twisting moments. In contrast, no correlation was observed in RPE1 cells, whose width/length ratios were between 0.5 and 0.8 (Figure 5A). A weak correlation was found in HeLa cells expressing PRC1-GFP, which had smaller width/length ratios than non-transfected HeLa cells (Figure S6A).

**Figure 5.**
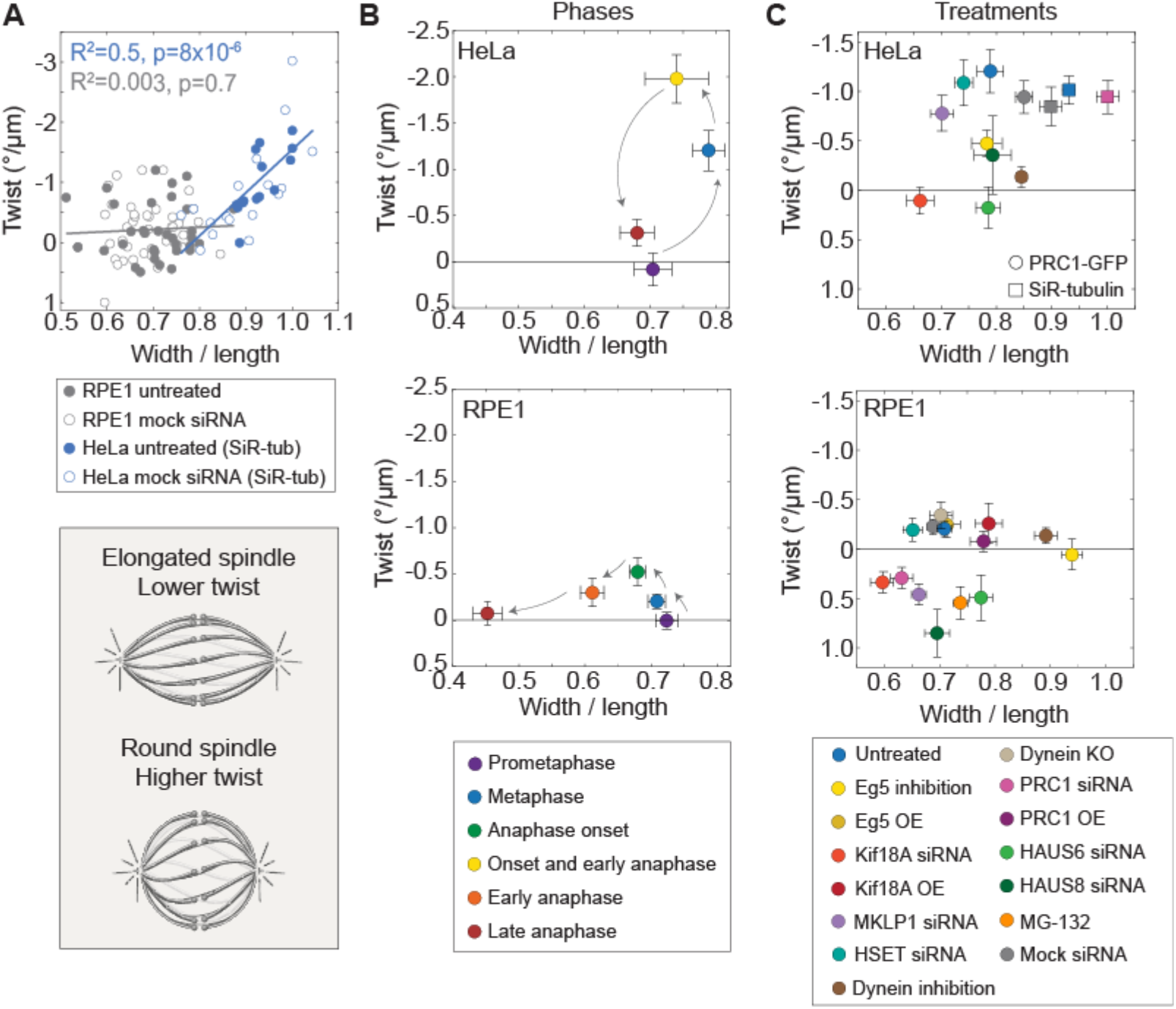
Round spindles have stronger twist than elongated spindles. **(A)** Graph shows the correlation between width/length ratio and twist in HeLa and RPE1 cells. Filled circles represent untreated cells while empty circles represent mock siRNA controls as indicated in the legend below the graph. Lines show linear fits for each cell line (untreated cells together with mock siRNA controls); equations y=-7.16x+5.62 for non-transfected HeLa cells and y=-0.36x+0.04 for RPE1 cells; goodness of fit is shown in the graph. Data for HeLa cells were also used in graphs in Figure 4B. Data for RPE1 cells were also used in graphs in Figures 1E, 3C, 4C. Experiments were performed on the nontransfected HeLa cell line and hTERT-RPE1 cells, permanently transfected and stabilized using CENP-A-GFP and centrin1-GFP. The scheme on the bottom shows that elongated and round spindles, with lower and higher width/length ratio, have lower and higher twist, respectively. **(B)** Graphs showing how the spindle twist and width/length ratio in HeLa (top) and RPE1 cells (bottom) vary over different phases of mitosis. Each color represents a phase, as described in the legend below the graphs; error bars represent SEM; arrows indicate progression of mitosis. The same data were used in graph in Figures 1D, 1E. Experiments were performed on the HeLa-Kyoto BAC cells stably expressing PRC1-GFP and hTERT-RPE1 cells permanently transfected and stabilized using CENP-A-GFP and centrin1-GFP. **(C)** Graphs showing how the spindle twist and width/length ratio in HeLa (top) and RPE1 cells (bottom) vary depending on the perturbations of spindle-associated proteins. Colors represent different protein perturbation, as described in the legend below the graphs; error bars represent SEM. The same data were used in graphs in Figures 3B, 3C, 4B, 4C. For HeLa cells, experiments were performed on HeLa-Kyoto BAC cells stably expressing PRC1-GFP (circles) and non-transfected HeLa cells stained with SiR-tubulin (rectangles), hTERT-RPE1 cells permanently transfected and stabilized using CENP-A-GFP and centrin1-GFP, and RPE1 inducible CRISPR/Cas9 DYNC1H1 knockout cells.

A plot of the twist as a function of the width/length ratio for various mitotic phases and treatments indicates that different combinations of twist and bending moments exist in spindles in different phases of mitosis or in which different molecular mechanisms are perturbed (Figures 5B, 5C; see Figure S6 for twist vs. width or length). In HeLa cells, prometaphase and late anaphase spindles are elongated with zero and small left-handed twist values, respectively (Figure 5B). Left-handed twist rises during metaphase when spindles are the roundest, and highest twist values are at the beginning of anaphase when spindles are still rather round (Figure 5B). In contrast, in RPE1 cells such correlation between twist and roundness over mitotic phases was not observed (Figure 5B).

When analyzing the twist of metaphase spindles across the treatments, we found that in HeLa cells, the left-handed twist was prevalent in spindles with mild or high width/length ratios (higher than ~0.8), whereas both left-handed and right-handed twist was found in spindles with lower width/length ratios (lower than ~0.8; Figures 5C). In RPE1 cells, left-handed twist was found in spindles over the whole range of width/length ratios, and had overall smaller values than in HeLa cells, whereas right-handed twist was found in spindles with lower width/length ratios (lower than ~0.8, Figures 5C). Taken together, these results suggest that strong left-handed twist can be found in rounder spindles, and right-handed twist in more elongated spindles.

## DISCUSSION

### Mechanisms that generate spindle twist

The chiral shape of the human mitotic spindle, visible as the left-handed twist of microtubule bundles, implies that torques act within the spindle. In this work we reveal biomechanical and molecular mechanisms that regulate the torques within microtubule bundles reflected in the spindle twist. From a biomechanical point of view, we show that forces within or outside the spindle regulate spindle twist (Figure 6A, box 1). Among the spindles in non-transfected HeLa cells during metaphase, round spindles are more twisted than elongated ones. In agreement with this, HeLa cell spindles in metaphase and just after anaphase onset are more round and more twisted than in prometaphase and late anaphase, when the spindles are elongated and twist is largely absent. In RPE1 spindles, which are overall more elongated than HeLa spindles, the twist is weaker and not correlated with the width/length ratio. Moreover, when we squeezed HeLa spindles along the pole-to-pole axis, they became rounder and their twist increased. These findings suggest that spindle roundness, which reflects bending moments within the spindle^28^, is correlated with twist. Thus, the molecular mechanisms that generate larger bending moments, causing the spindles to be rounder, may also generate larger twisting moments, visible as stronger twist of the microtubule bundles. It is interesting to see that spindles, as complex and dynamic structures, show a relationship between twisting and bending similar to simple systems from classical beam mechanics^59^.

**Figure 6.**
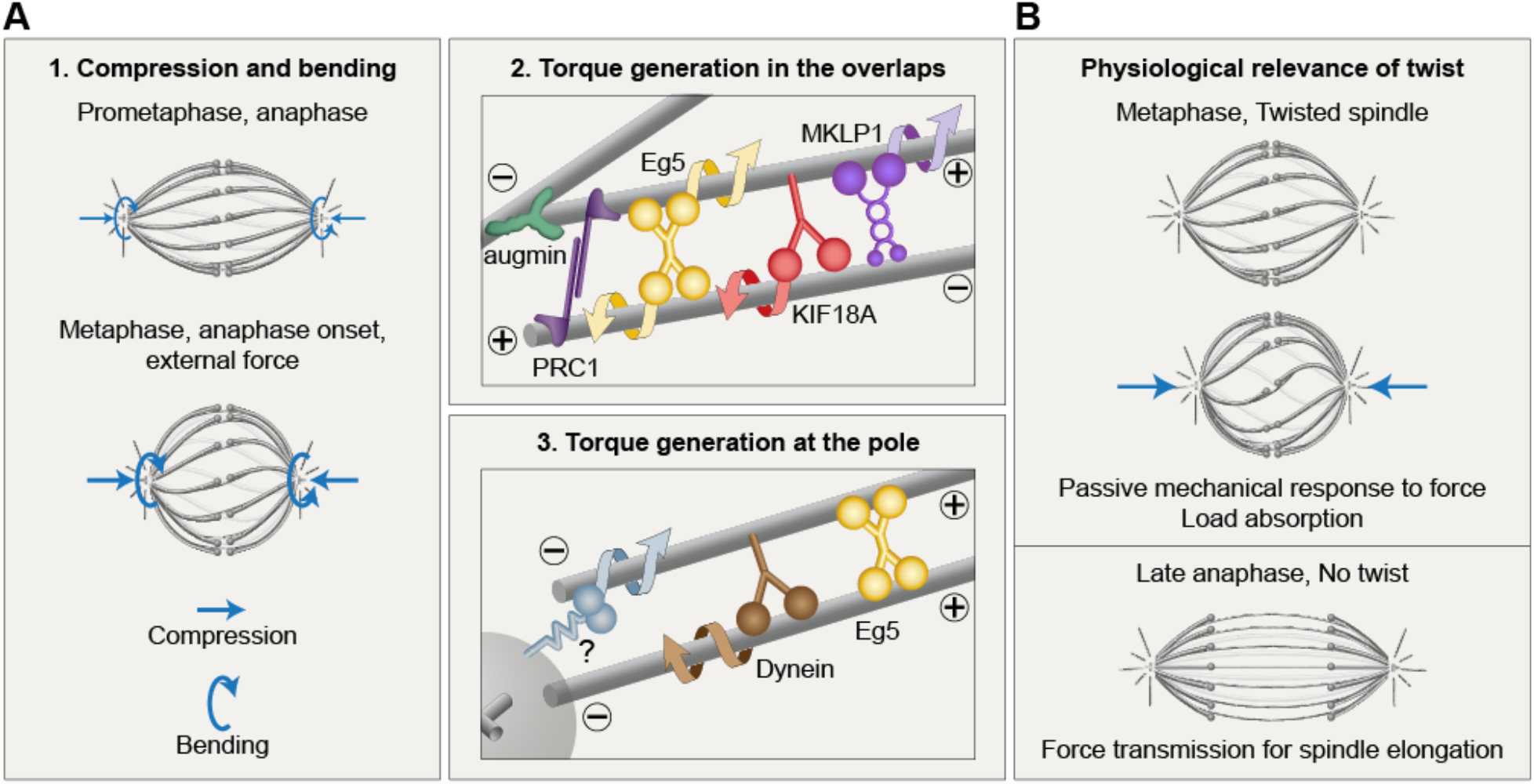
Biomechanical and molecular origins of spindle twist and its biological role. **(A)** Forces regulate twist (box 1). Round spindles or those compressed by external forces (blue straight arrows) are more twisted than elongated ones, suggesting that larger bending moments (blue curved arrows) are correlated with larger twist. Within the antiparallel overlaps of bridging microtubules (box 2), Eg5, Kif18A, and MKLP1 rotate the microtubules around one another, whereas crosslinking by PRC1 constrains the free rotation of microtubules within the bundle, allowing for accumulation of torsional stresses. Augmin contributes to the twist by nucleating bridging microtubules. At the spindle pole (box 3), Eg5 crosslinks parallel microtubules, which may prevent their free rotation. Eg5 and other motors (question mark) may rotate the microtubules around the pole. Dynein contributes to torque generation by moving in a minus-end directed helical manner. **(B)** Spindle twist allows for a mechanical response to external forces by absorbing load during metaphase (top). In contrast, in late anaphase twist is absent, which promotes force transmission for spindle elongation and maintenance of chromosome separation (bottom).

By performing a candidate screen in which we perturbed motor proteins that step in a chiral manner and other microtubule-associated proteins, we identified several molecular players involved in the regulation of spindle chirality. Inactivation of Eg5 decreased the left-handed twist in HeLa cells, in agreement with our previous findings^28^, and led to zero twist in RPE1 cells. Depletion of Kif18A resulted in zero twist in HeLa cells and a switch of the twist direction from left-handed to right-handed in RPE1 cells. Overexpression of these proteins did not change the twist. Depletion of MKLP1 in RPE1 cells also resulted in a change of twist to the right-handed direction. All three motors are known to exert torque on the microtubules *in vitro*^34,36,40,42^, and are found within the antiparallel overlaps of bridging microtubules in the spindle^12,54,60^. Thus, we suggest that they generate the twisted shape of the bundle by rotating the antiparallel microtubules within the bundle around each other (Figure 6A, box 2). Eg5 may also contribute to spindle twist by acting at the spindle pole (Figure 6A, box 3), given that the most pronounced localization of Eg5 is in the pole region^12,60^. Here, Eg5 may crosslink parallel microtubules^61,62^, which would prevent their free rotation within the bundle and promote the accumulation of torsional stresses, but it may also actively generate torques. Additionally, other motors localized at the pole, such as Kif2a^63^ and Kif2c/MCAK^64^ from the kinesin-13 family, may contribute to spindle twist by rotating the microtubules with respect to the spindle pole (Figure 6A, box 3). In HeLa cells, inhibition of dynein abolished the spindle twist. Cytoplasmic dynein shows a bidirectional helical motility that can generate torque in either direction, but prefers right-directed movement^37^. Thus, its net torque generation may also influence spindle twist (Figure 6A, box 3).

Intriguingly, the strongest effect on spindle twist was obtained by depletion of the HAUS6 and HAUS8 subunits of the augmin complex, which promotes nucleation of new microtubules from the wall of existing microtubules^48,55,65^. These depletions resulted in zero twist in HeLa cells and right-handed twist in RPE1 cells. Given that augmin depletion reduces the microtubule number within bridging fibers^56^, we suggest that the altered twist is related to the reduced antiparallel overlaps where torque-generating motors such as Eg5, Kif18A, and MKLP1 bind (Figure 6A, box 2). In addition, augmin nucleates new microtubules at an angle with respect to the wall of the old microtubule^66,67^, thus it is tempting to speculate that if the new microtubules extend skewed from the old microtubules and spiral around them, this may lead to the twist of the entire microtubule bundle.

Depletion or overexpression of the crosslinker of antiparallel microtubules PRC1 resulted in no twist in RPE1 cells. Though the effects were subtle, we speculate that in a metaphase spindle the microtubules crosslinked by PRC1 cannot rotate freely within the bundle (Figure 6A, box 2), resulting in the accumulation of torsional stresses, yet the torsional rigidity of the whole bundle is low enough to allow for twisting of the bundle. In contrast, in late anaphase when PRC1 is highly abundant on midzone microtubules, or upon PRC1 overexpression in metaphase, the increased number of PRC1-bundled microtubules may increase the torsional rigidity of the entire bundle, making it harder to twist.

Duration of metaphase also affects the spindle twist. Arresting RPE1 cells in metaphase resulted in right-handed twist, similar to the depletion of Kif18A or the augmin complex. Interestingly, these depletions also prolong metaphase^47,48^, which may contribute to the observed effect on twist.

The switch of the direction of twist from left-handed to right-handed indicates the existence of competing mechanisms promoting twist in the opposite directions. Different perturbations may tip the balance one way or the other. HeLa spindles have stronger left-handed twist than RPE1 spindles, and protein depletions that led to zero twist in HeLa cells largely resulted in right-handed twist in RPE1 cells, thus the twist changed in both cell lines by a similar amount. This implies that torques are regulated by similar mechanisms in both cell lines, but the torque balance is shifted more towards the formation of left-handed twist in HeLa than in RPE1 spindles. Unlike metaphase, in anaphase twist is mostly absent in HeLa and RPE1 spindles. A recent study found strong left-handed twist during anaphase after combined Eg5 inhibition and NuMa depletion in RPE1 cells, suggesting that opposing motors are required to prevent twisting in the anaphase spindle^29^.

All of the diverse molecular perturbations used in this work tuned the twist in the same direction, towards more positive values, suggesting that the corresponding molecular players promote left-handed twist. We thus speculate that right-handed twist may arise due to the activity of microtubule-associated proteins that were not covered by our candidate screen. Additionally, the helical structure of the microtubule lattice^68^ may influence the twist of a bundle of microtubules.

In contrast to the left-handed twist of human spindles, spindles in the amoeba *Naegleria gruberi* are twisted in a right-handed fashion^30^, which may be due to the differences in kinesins and other microtubule-associated proteins between *Naegleria* and humans. *Naegleria* lacks homologs to subunits of the augmin complex^69^, which is in line with the right-handed twist of spindles in this amoeba and in RPE1 cells depleted of augmin. Helical structures are also widespread in plants, e.g., *lefty* mutants in *Arabidopsis thaliana* have cortical microtubule arrays that form right-handed helices, resulting in clockwise bending of leaf petioles and flower petals when viewed from above^70^, whereas *spiral* mutants show counterclockwise bending^*71*^. What determines the direction and amount of twist in microtubules of different organisms and whether there are common elements remains an intriguing topic for future studies.

### Physiological function of spindle twist

Although spindle chirality may be simply a side-effect of the action of motors that generate torque, the twisted shapes of microtubule bundles may contribute to spindle function. We propose that the twisted shape observed during metaphase is beneficial for the spindle because it allows for changes of spindle shape as a mechanical response to external forces. In this picture, a twisted spindle can quickly shorten under compressive forces by increasing the twist in a manner similar to an elastic spring (Figure 6B, top). This response does not require depolymerization of microtubules during spindle shortening, which would be the case for non-twisted spindles. Our experiments in which we compressed the spindle along the pole-to-pole axis and observed an increase in twist as the spindle shortened, while the contour length of microtubule bundles remained largely unchanged, provide support for the model in which the built-in twist helps the spindle to respond to forces and decreases the risk of spindle breakage under high load.

In contrast to metaphase, during late anaphase, the spindle is not chiral as the bundles lose their twist and become straight. This straightening is likely due to the accumulation of PRC1 and also other midzone proteins within these bundles. We speculate that the straight shapes of the bundles are beneficial for the spindle in late anaphase to allow for force transmission from the central overlap region to the poles, to move the chromosomes apart and keep them separated (Figure 6B, bottom).

Additional functions of spindle chirality may be to promote physical separation of adjacent bundles during prometaphase or to help start spindle elongation at the onset of anaphase by releasing elastic energy stored in the twisted bundles. Intriguingly, a recent study showed that changes in twist can be associated with chromosome segregation errors^29^. Thus, regulation of twist may be important for the fidelity of chromosome segregation, which awaits further explorations. Overall, we expect that the results presented here will motivate exciting new research on the molecular mechanisms and the biological roles of rotational forces in the spindle.

## MATERIALS AND METHODS

### Cell lines

Experiments were performed using: HeLa-Kyoto BAC cell line stably expressing PRC1-GFP courtesy of Ina Poser and Tony Hyman (Max Planck Institute of Molecular Cell Biology and Genetics, Dresden, Germany); unlabeled (non-transfected) HeLa-TDS cells from the High-Throughput Technology Development Studio (MPI-CBG, Dresden); human hTERT-RPE1 permanently transfected and stabilized using CENP-A-GFP and centrin1-GFP courtesy of Alexey Khodjakov (Wadsworth Center, New York State Department of Health, Albany, NY); inducible CRISPR/Cas9 DYNC1H1 knockout RPE1 cells line courtesy of Iain M. Cheeseman (Massachusetts Institute of Technology, Cambridge, MA, USA). Cells were grown in flasks in Dulbecco’s modified Eagle’s medium (DMEM; Capricorn Scientific GmbH, Germany) supplemented with 10% fetal bovine serum (FBS; Sigma-Aldrich, MO, USA), 10000 U/ml penicillin/streptomycin solution (Capricorn Scientific GmbH, Germany), and for HeLa-Kyoto BAC cells also 50 μg/mL geneticin (Life Technologies, Waltham, MA, USA). The cells were kept at 37 °C and 5% CO2 in a Galaxy 170S CO2 humidified incubator (Eppendorf, Hamburg, Germany) and regularly passaged at the confluence of 70-80%.

### Sample preparation

To visualize microtubules in experiments on RPE1 and non-transfected HeLa cells, silicon rhodamine (SiR)-tubulin (λ_Abs_ 652 nm, λ_Em_ 674 nm) (Spirochrome AG, Stein am Rhein, Switzerland) dye was added to the dish at the final concentration of 100 nM, 2-3 hours prior to imaging. To visualize chromosomes and determine phase of the mitosis of the spindle in experiments on HeLa PRC1-GFP cells, 1 hour prior to imaging silicon rhodamine (SiR)-DNA (λ_Abs_ 652 nm, λ_Em_ 674 nm) (Spirochrome AG, Stein am Rhein, Switzerland) was added to the dish at a final concentration of 100 nM. To visualize chromosomes and determine phase of the mitosis of the spindles in experiments on non-transfected HeLa cells and RPE1 inducible DYNC1H1 knockout cells, 50 μL of NucBlue Live Ready Probes Reagent (Hoechst 33342) (Invitrogen by Thermo Fischer Scientific, MA, USA) dye was added to the dishes, 1 min before imaging. Lipofectamine RNAiMAX reagent (Invitrogen by Thermo Fisher Scientific, MA, USA) was used for RNAi treatments following manufacturer’s instructions. Transfections with siRNA were always performed 48 hours prior to imaging at the final concentration of 100 nM. For the inhibition of Eg5, cells were treated with (+)-S-Trityl-L-cysteine (STLC, Sigma-Aldrich, MO, USA) at the final concentration of 40 μM right before the imaging so that cells are not yet collapsed into a monopol during imaging. STLC-treated cells were imaged before spindle shortening (up to 5 min in STLC) and after shortening (10-20 min in STLC). For the inhibition of dynein, cells were treated with dynarrestin (HY-121802/CS-0083323, MedChemExpress, NJ, USA) at the final concentration of 50 μM, 1 hour prior to imaging, and were imaged up to 2 hours after the addition of the drug. This time period allowed the spindles to shorten, which was used to confirm that the inhibition experiment worked^51^. CRISPR/Cas9 knockout of DYNC1H1 in RPE1 cell line was induced with doxycycline hyclate (D9891-1G, Sigma-Aldrich, MO, USA) at the final concentration of 1 μg/mL at 24 hour intervals for 4 consecutive days, with imaging and analysis on the fifth day. Here, only spindles with splayed poles, but bipolar shape were imaged. For depletion of endogenous Kif18A, cells were transfected with Kif18A Silencer Select siRNA (4390825, Ambion, Thermo Fisher Scientific, MA, USA). For depletion of endogenous PRC1, cells were transfected with ON-TARGETplus SMARTpool Human PRC1 (L-C19491-00-0010, Dharmacon, CO, USA). For depletions of endogenous HAUS6 and HAUS8, cells were transfected with ON-TARGETplus SMARTpool Human HAUS6 (L-018372-01-0005, Dharmacon, CO, USA) and ON-TARGETplus SMARTpool Human HAUS8 (L-031247-01-0005, Dharmacon, CO, USA), respectively. For the depletion of endogenous HSET, cells were transfected with ON-TARGETplus SMART pool Human KIFC1 (L-004958-00, Dharmacon, CO, USA). For the depletion of endogenous MKLP1, cells were transfected with siRNA (sc-35936; Santa Cruz Biotechnology, TX, USA). In mock experiments cells were transfected with equal amount of ON-TARGETplus Control Pool Non-Targeting pool (D-001810-10-05, Dharmacon, CO, USA) or Silencer Select Negative Control #1 siRNA (4390843, Ambion, Thermo Fisher Scientific, MA, USA). All plasmid transfections were performed using Nucleofactor Kit R with the Nucleofactor 2b Device (Lonza, Basel, Switzerland) using Y-001 program for human HMEC cells (high efficiency). To overexpress Eg5 protein, cells were transfected with 5 μg of mEmerald-Kinesin11-N-18 plasmid (Addgene number: 54137) 24 hours prior to imaging. For Kif18A overexpression, cells were transfected with 5 μm of EGFP-Kif18A plasmid that was a gift from Jason Stumpff (University of Vermont, Burlington, VT, USA). To overexpress PRC1 protein, cells were transfected with 5 μg of mCherry-PRC1 plasmid that was a gift from Casper C. Hoogenraad (Utrecht University, Utrecht, Netherlands). To reduce the time in which HeLa cells reach anaphase, the Mps1 inhibitor AZ3146 (Sigma-Aldrich, MO, USA) was added in prometaphase at a final concentration of 4 μM. Metaphase arrest in RPE1 cells was performed with the proteasome inhibitor MG-132 (M7449, Sigma-Aldrich, MO, USA) added at least 2 hours prior to imaging at a final concentration of 20 μM. All experiments were performed at least three times in both cell lines, except Kif18A overexpression that was preformed once. To prepare samples for microscopy, RPE1 and HeLa cells were seeded and cultured in DMEM medium with supplements at 37 °C and 5% CO2 on uncoated 35-mm glass coverslip dishes with 0.17-mm (1.5 coverglass) glass thickness (MatTek Corporation, Ashland, MA, USA).

### Immunofluorescence

HeLa-Kyoto BAC cell line stably expressing PRC1-GFP were grown on glass-bottomed dishes (14 mm, No. 1.5, MatTek Corporation) and fixed by a microtubule-preserving mixture of 3.2% PFA (paraformaldehyde) and 0.25% GA (glutaraldehyde) in microtubule-stabilizing PEM buffer (0.1 M PIPES, 0.001 M MgCl_2_ x 6 H_2_O, 0.001 M EDTA, 0.5% Triton-X-100) for 10 min at room temperature. After fixation with PFA and GA, for quenching, cells were incubated in 1 mL of freshly prepared 0.1% borohydride in PBS (phosphate-buffered saline) for 7 min and after that in 1 mL of 100 mM NH_4_Cl and 100 mM glycine in PBS for 10 min at room temperature. Cells were then washed with 1 mL of PBS, 3 times for 5 min. To block unspecific binding of antibodies, cells were incubated in 500 μL blocking/permeabilization buffer (2% normal goat serum (NGS) and 0.5% Triton-X-100 in water) for 45 min at room temperature. Cells were then incubated in 500 μL of primary antibody solution (rat anti-alpha Tubulin YL1/2 (MA1-80017, Invitrogen, CA, SAD), diluted 1:500) for 24 hours at 4 °C. After primary antibody, cells were washed in PBS and then incubated in 500 μL of secondary antibody solution (donkey anti-rat IgG Alexa Fluor 594 (ab150156, Abcam), diluted 1:1000) for 45 min at room temperature.

Human hTERT-RPE1 cells, permanently transfected and stabilized using CENP-A-GFP and centrin1-GFP, were grown on glass-bottomed dishes (as described above) and fixed in cold 100% methanol for 1 min on the ice block. After fixation, cells were washed in PBS 3 times for 5 min at room temperature. Next, cells were additionally permeabilized in 0.5% Triton-X-100 solution for 15 min at room temperature and then washed in PBS (as described above). To block unspecific binding of antibodies, cells were incubated in 1% NGS solution for 1 hour on 4 °C. After washing in PBS once, cells were incubated with primary antibodies (dilution 1:100 in 1% NGS) overnight on 4 °C. Next, cells were washed in PBS 3 times for 5 min at room temperature and incubated with secondary antibodies (dilution 1:250 in 2% NGS solution) for 1 hour at room temperature covered with aluminum foil. Before microscopy, cells were washed in PBS (as described above) and left in PBS during imaging. Cells were kept in the dark in PBS on 4 °C. Primary antibodies used: PRC1 (C-1) mouse monoclonal IgG_1_ (sc-376983, Santa Cruz Biotechnology, TX, USA), Rabbit anti-KIF18A Affinity Purified (A301-080A, Bethyl, TX, USA), Rb pAb to FAM29A (ab150806, Abcam, Cambridge, UK), HICE1 Polyclonal Antibody (PA5-21331, Invitrogen, MA, USA), KIFC1 (M-63) mouse monoclonal IgG_2a_ (sc-100947, Santa Cruz Biotechnology, TX, USA), Rb pAb to MKLP1 (ab174304, Abcam, Cambridge, UK); secondary antibodies used: Dnk pAb to Ms IgG (ab150112, Abcam, Cambridge, UK), Dnk pAb to Rb IgG (ab150064, Abcam, Cambridge, UK).

### Spindle compression

Spindle compression method was optimized from Mitchison and Dumont, 2009 ^46^. A solution of 2% ultra-pure agarose (15510 Invitrogen by Thermo Fisher Scientific, MA, USA) in PBS was prepared, brought to boil and 2 mL was put in a 35 mm petri dish to solidify with ~2 mm thickness. A 1 cm × 1 cm pad area was cut out, soaked in L-15 medium overnight at 4 °C for equilibration, and warmed to 37 °C just before use. Cells were plated on 14 or 20 mm glass microwell uncoated dishes before imaging. A flat metaphase cell was chosen among 80-100% confluent cells for pre-perturbation imaging. After imaging of the metaphase cell before compression, the pad was deposited gently, centered on the cell. Note: it is important to do this step gently and with minimal moving of the dish so the position of the cell could stay intact. Using an oil hydraulic fine manipulator (InjectMan 4, micromanipulator with dynamic movement control, 100–240 V/50–60 Hz) and a coarse manipulator attached to the confocal microscope. A metal rod (which is a part of micromanipulator where the needle for microinjection is inserted) was centered on the cell and lowered (z-axis) until weak contact was made with the pad (rod diameter ≫ cell diameter). The rod was lowered slowly (over ~10 s) for several μm until the cell area expanded, and its position kept constant as the cell and spindle responses were imaged. HeLa PRC1-GFP cells were imaged every 30 s for 3 times which gave us 4 different times the cell was imaged at: before compression, 0.5 min after compression, 1 min after compression and 1.5 min after compression. Cell health was monitored through the presence of the intact cell membrane and the ability of the cell to enter anaphase after perturbation. Rough estimate of the number of vertically oriented metaphase spindles in a 35 mm petri dish is around 1 spindle per 50-100 horizontally oriented metaphase spindles. From 31 spindles that were compressed in about 45 independent experiments, 23 spindles rotated during the first 30 s of compression and could not be used for further analysis. From 8 spindles that remained vertically oriented during compression, 2 spindles were not compressed enough which was determined by their length. Successful compression, where spindle length decreased for 0.5 μm or more, was achieved on 6 spindles from 5 independent experiments, which were then used for the analysis.

### Confocal microscopy

Live RPE1 and HeLa cells were imaged using Bruker Opterra Multipoint Scanning Confocal Microscope^72^ (Bruker Nano Surfaces, Middleton, WI, USA). The system was mounted on a Nikon Ti-E inverted microscope equipped with a Nikon CFI Plan Apo VC ×100/1.4 numerical aperture oil objective (Nikon, Tokyo, Japan). During imaging, cells were maintained at 37 °C in Okolab Cage Incubator (Okolab, Pozzuoli, NA, Italy). A 22 μm slit aperture was used for RPE1 and 60 μm pinhole for HeLa cells. The xy-pixel size was 83 nm. For excitation of GFP and mCherry fluorescence, a 488 and a 561 nm diode laser line was used, respectively. For SiR-dyes, a 640 nm diode laser line was used. The excitation light was separated from the emitted fluorescence by using Opterra Dichroic and Barrier Filter Set 405/488/561/640. Images were captured with an Evolve 512 Delta EMCCD Camera (Photometrics, Tucson, AZ, USA) with no binning performed. To cover the whole metaphase spindle, z-stacks were acquired at 30–60 focal planes separated by 0.5 μm with unidirectional xyz scan mode. The system was controlled with the Prairie View Imaging Software (Bruker Nano Surfaces, Middleton, WI, USA).

### Image and data analysis

To calculate spindle twist, microscopy images of horizontal spindles were analyzed in Fiji Software (ImageJ, National Institutes of Health, Bethesda, MD, USA)^73^. Only images with both spindle poles in the same plane before and during imaging of the z-stack were used in analysis to avoid unspecific spindle movements in the calculation of spindle twist. Horizontal spindles were transformed into vertical orientation using a code written in R programming language in RStudio^28^. In transformed stack microtubule bundles and poles appear as blobs.

#### Visual assessment

In this method, the spindle is observed end-on and the rotation of microtubule bundles around the pole-to-pole axis is estimated visually. If the bundles rotate clockwise when moving along the spindle axis in the direction towards the observer, the twist is left-handed, and vice versa (Figure S1A, left). The outcome of our visual assessment is a score of spindle twist, which describes whether the spindle has a left-handed, weak left-handed, righthanded, weak right-handed, or no visible twist. Weak left-handed or weak right-handed twists correspond to a range of approximately −1 to −2 °/μm in the bundle tracing method. This is visible as a total rotation of 5-10° in the clockwise (left-handed) or counter-clockwise (right-handed) direction in the end-on view of the spindle when moving towards the observer along the bundle length, where bundles are typically 5 μm long. Left-handed or right-handed twists correspond to a rotation of more than 10° in the end-on view. The advantage of this method is its trustworthiness because coarse classification of spindles into 5 groups is reliable, whereas the main disadvantage is that the results are semi-quantitative rather than quantitative.

#### Optical flow

In the optical flow method, the movement of the signal coming from microtubule bundles is estimated automatically by comparing the signal from one z-plane to the next (Figure S1A, middle). This method yields a value for the average twist of all bundles in a spindle. It is the preferred choice for experiments on a large number of spindles because it is automated. Disadvantages are that it provides only the average twist value rather than the twist of each bundle, and that the results are sensitive to unspecific signal in the images, individual bundles with atypical behavior, and imperfect alignment of the spindle axis with the z-axis.

First, parts of the images containing the blobs were selected for analysis using Rectangle tool in ImageJ. In all transformed stacks only images between spindle poles were used for analysis. Transformed spindle images contained a lot of noise that was removed by using the Mexican hat filter and a threshold. The Mexican hat filter, also called the LoG (Laplacian of Gaussian) filter, was used for detection of blobs^74,75^. After applying the Mexican hat filter, a threshold was applied to the image. It removes all the pixels with intensity lower than the given threshold. Microtubule bundles of transformed spindles were detected and traced automatically using optical flow for calculating the movement of pixels between two consecutive images. Farnebäck’s two-frame motion estimation algorithm (dense optical flow algorithm) was used ^76^. The spindle poles were tracked manually using Multipoint tool in ImageJ. Helicities of spindles were calculated using the algorithm called All pixels weighted helicity algorithm. It calculates the total helicity as the average helicity of all pixels in the spindle, weighted by their normalized intensity. The tilt of the spindle with regard to the imaging plane was calculated from the tracked spindle poles, and the twist measurement was corrected by this tilt angle. The code for tracing of bundles and helicity calculating was written in Python programming languange using PyCharm IDE. The external libraries used in image preprocessing, calculating helicity and visualisation are NumPy, scikit-image, Matplotib, PIL, OpenCV and SciPy. The code and instructions are available at https://gitlab.com/IBarisic/detecting-microtubules-helicity-in-microscopic-3d-images.

#### Bundle tracing

Bundles in images of spindles oriented vertically were traced manually using Multipoint tool in Fiji^28^. We convert the imaging plane (z-plane) to its corresponding z-coordinate by multiplying with the distance between successive planes set during image acquisition (0.5 μm) and by a factor of 0.81 to correct for the refractive index mismatch^28^. Next, to describe the shape of a microtubule bundle, we use the Oblique circle method^43^. We first position the spindle so that the pole-to-pole axis is aligned with our coordinate system, i.e., we untilt the spindle. Next, we fit a plane to the points representing the bundle, and then we fit a circle that lies in this plane to the same points. From these fits we calculate the curvature and twist of the bundle as follows: (i) The curvature is calculated as one over the radius, and (ii) the twist is calculated as the angle between the plane and the z-axis divided by the mean distance of these points from the z-axis. Contour length of the bundle was calculated as the length of the fitted circular arc plus the distance of bundle ends from the corresponding poles. The main advantage of this method is that it yields a value of twist for each individual bundle in the spindle, whereas the main disadvantage is that it requires manual tracing, which makes it impractical for high-throughput studies.

To measure spindle length and width, we used the Line tool in Fiji. Length was measured by drawing a line from pole to pole of the spindle. Width in HeLa cells expressing PRC1-GFP was measured by drawing a line across the equatorial plane of the spindle, with the line ending at the outer edges of a spindle. Width in RPE1 cells expressing CENP-A-GFP and centrin1-GFP was measured by drawing a line across the equatorial plane of the spindle, with the line ending at the outer kinetochore pairs.

To quantify protein expression, the fluorescence intensity signal of the protein of interest was measured on the whole spindle region using ImageJ Polygon Selection tool on the sumintensity projection of the whole z-stack. The mean background fluorescence intensity measured in the cytoplasm was subtracted from the mean value obtained on the spindle, and the resulting value was divided by the number of z-slices used in the sum projection.

### Statistical analysis

Fiji was used to scale images and adjust brightness and contrast. Figures were assembled in Adobe Illustrator CS5 and CC (Adobe Systems, Mountain View, CA, USA). Graphs were plotted in MATLAB (MathWorks, Natick, MA, USA). For generation of univariate scatter plots, the open “UnivarScatter” Matlab extension was used (https://github.com/manulera/UnivarScatter). Data are given as mean ± s.e.m., unless otherwise stated. Significance of data was estimated by Student’s t-test (two-tailed and two sample unequal-variance; except for the experiments with spindle compression, where a paired t-test was used to compare the values for the same spindles before and after compression). p < 0.05 was considered statistically significant. Values of all significant differences are given with degree of significance indicated (*0.01 <p < 0.05, **0.001 < p < 0.01, ***p < 0.001). Statistically significant differences between groups of data were determined by one-way ANOVA and Tukey’s HSD post hoc test, p < 0.05 was considered statistically significant. The number of analyzed cells and microtubule bundles is given in the respective figure panel.

## ACKNOWLEDGEMENTS

We thank Ina Poser, Tony Hyman, Alexey Khodjakov, and Iain M. Cheeseman for cell lines; Jason Stumpff and Casper C. Hoogenraad for plasmids; Juraj Simunić, Jurica Matković, Valentina Štimac, Isabella Koprivec, Lucija Bujanić, and Iva Buljan for their contribution to experimental work; Martina Manenica for help with the MATLAB codes, and Ivana Šarić for the drawings. We thank Nenad Pavin, Maja Novak, Josip Tambača, Matko Ljulj, and all members of Tolić and Pavin groups for helpful discussions. This work was funded by the European Research Council (ERC Consolidator Grant, GA Number 647077), the Croatian Science Foundation (HRZZ project IP-2019-04-5967), the Science and Innovation Grant co-financed by the European Structural and Investment Funds (ESIF) within the Operational Programme Competitiveness and Cohesion (OPCC) 2014–2020 (Grant KK.01.1.1.04.0057), and the QuantiXLie Center of Excellence, a project co-financed by the Croatian Government and European Union through the European Regional Development Fund—the Competitiveness and Cohesion Operational Programme (Grant KK.01.1.1.01.0004). We also acknowledge new support by the ERC (Synergy Grant, GA Number 855158). The work of doctoral students M.T. and A.I. have been supported in part by the “Young researchers’ career development project – training of doctoral students” of the Croatian Science Foundation.

## AUTHOR CONTRIBUTIONS

M.T. performed experiments on RPE1 cells, B.K. and M.T. performed experiments on HeLa cells, except compression experiments, which were done by I.P. M.T., B.K., and I.P. analyzed the experimental data. I.B. and S.Š. developed the optical flow method for twist measurement. A.I. measured the twist from bundle traces. M.T. and I.M.T. wrote the manuscript, with input from I.P., B.K., A.I., and other authors. M.T. assembled the figures. I.M.T. conceived the project and supervised the experiments.

## DECLARATION OF INTERESTS

The authors declare no competing interests.

## SUPPLEMENTARY INFORMATION

**Figure S1.**
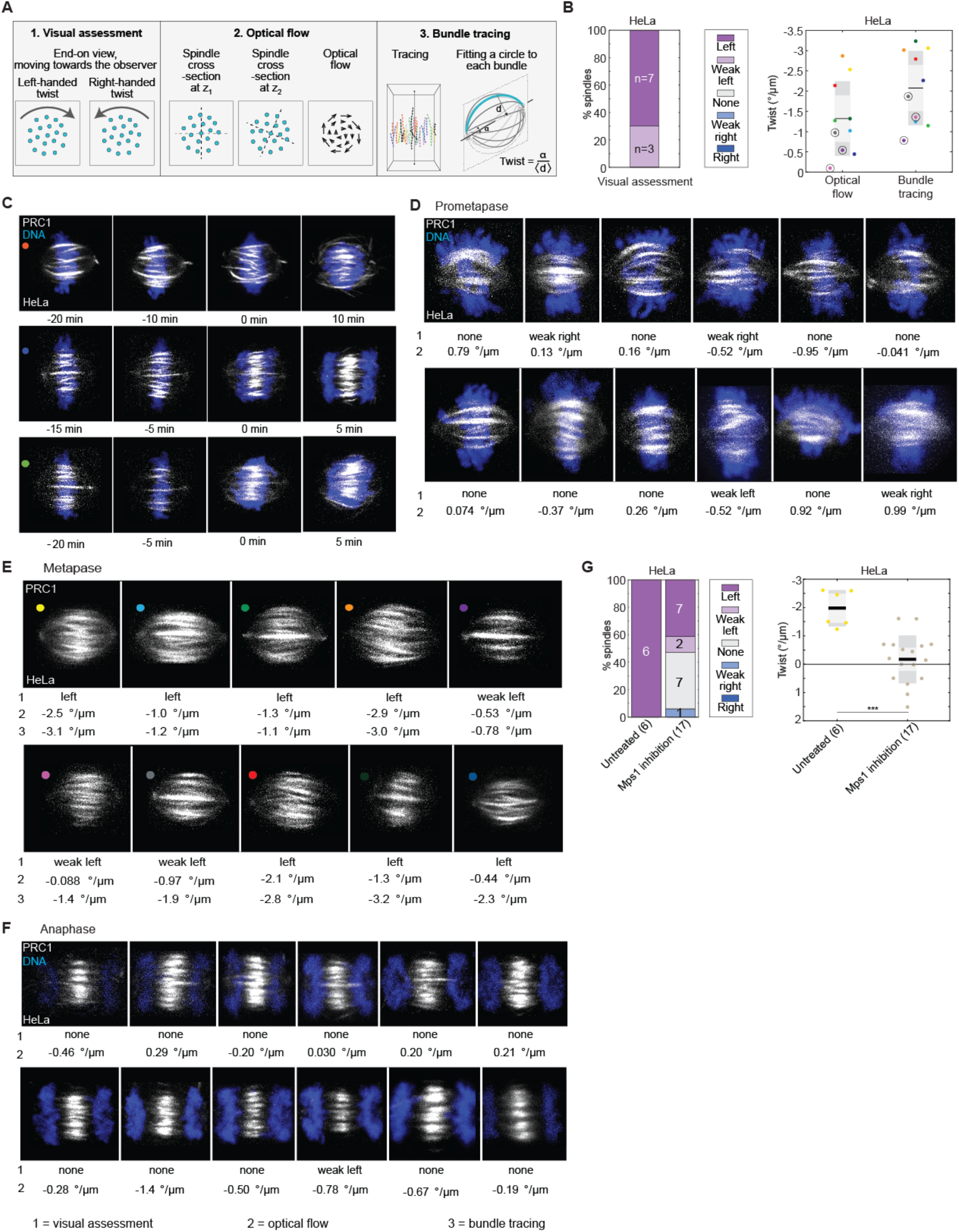
Methods for calculating twist and progression of twist during mitosis in HeLa cells. **(A)** Schemes of three methods used to measure spindle twist: visual assessment (1), optical flow (2) and bundle tracing (3). **(B)** Comparison of the twist for 10 spindles calculated with three different methods. On the left, visual assessment graph represents percentages of spindles showing left, right, weak left, weak right or no twist as described in the legend. On the right, graph shows twist calculated with optical flow and bundle tracing methods; each color represents one cell; circled and un-circled data correspond to the ‘weak left’ and ‘left’ data from the visual assessment graph, respectively. Note that the weak left twist in the visual assessment graph corresponds to the range of approximately −1 to −2 °/μm in the bundle tracing method followed along the 5 μm of the bundle length, which corresponds to the rotation of 5-10° in the clockwise direction in the end-on view of the spindle. The black line shows the mean; the light and dark grey areas mark 95% confidence interval on the mean and standard deviation, respectively. Same cells were used to calculate the data for both methods. Experiments were performed on the HeLa-Kyoto BAC cells stably expressing PRC1-GFP (n=10; raw data taken and re-calculated from ^28^). **(C)** Microscope images of HeLa cells’ spindles in time. Each colored dot represents one spindle’s progression through mitosis in time, and each color matches the color of the spindle’s data in the graph in Fig. 1C. Three individual examples are shown. Microtubule bundles are shown in grey (PRC1-GFP) and DNA in blue (SiR-DNA dye). Images are shown in maximum z-projections. Experiments were performed on the HeLa-Kyoto BAC cells stably expressing PRC1-GFP. **(D-F)** Microscope images of individual spindles of HeLa cells in different phases of mitosis. Examples of spindles in prometaphase (D), metaphase (E) and anaphase (F) are shown with their twist values. Twist was determined with the visual assessment method and the optical flow, marked 1 and 2, respectively. For metaphase spindles twist was also calculated using bundle tracing method, marked 3. Data from these cells was used in the graphs in Fig. 1D. Data from metaphase cells was also used in Fig. S1B. Microtubule bundles are shown in grey (PRC1-GFP) and DNA in blue (SiR-DNA dye). Images are shown in maximum z-projections. Experiments were performed on the HeLa-Kyoto BAC cells stably expressing PRC1-GFP. **(G)** Graphs showing twist values after the inhibition of Mps1 in HeLa cells. On the left, visual assessment graph represents percentages of spindles showing left, right, weak left, weak right or no twist as described in the legend. On the right, graph shows twist calculated with the optical flow method. The black line shows the mean; the light and dark grey areas mark 95% confidence interval on the mean and standard deviation, respectively. ***p<0.001 (Student’s t-test). Experiments were performed on the HeLa-Kyoto BAC cells stably expressing PRC1-GFP.

**Figure S2.**
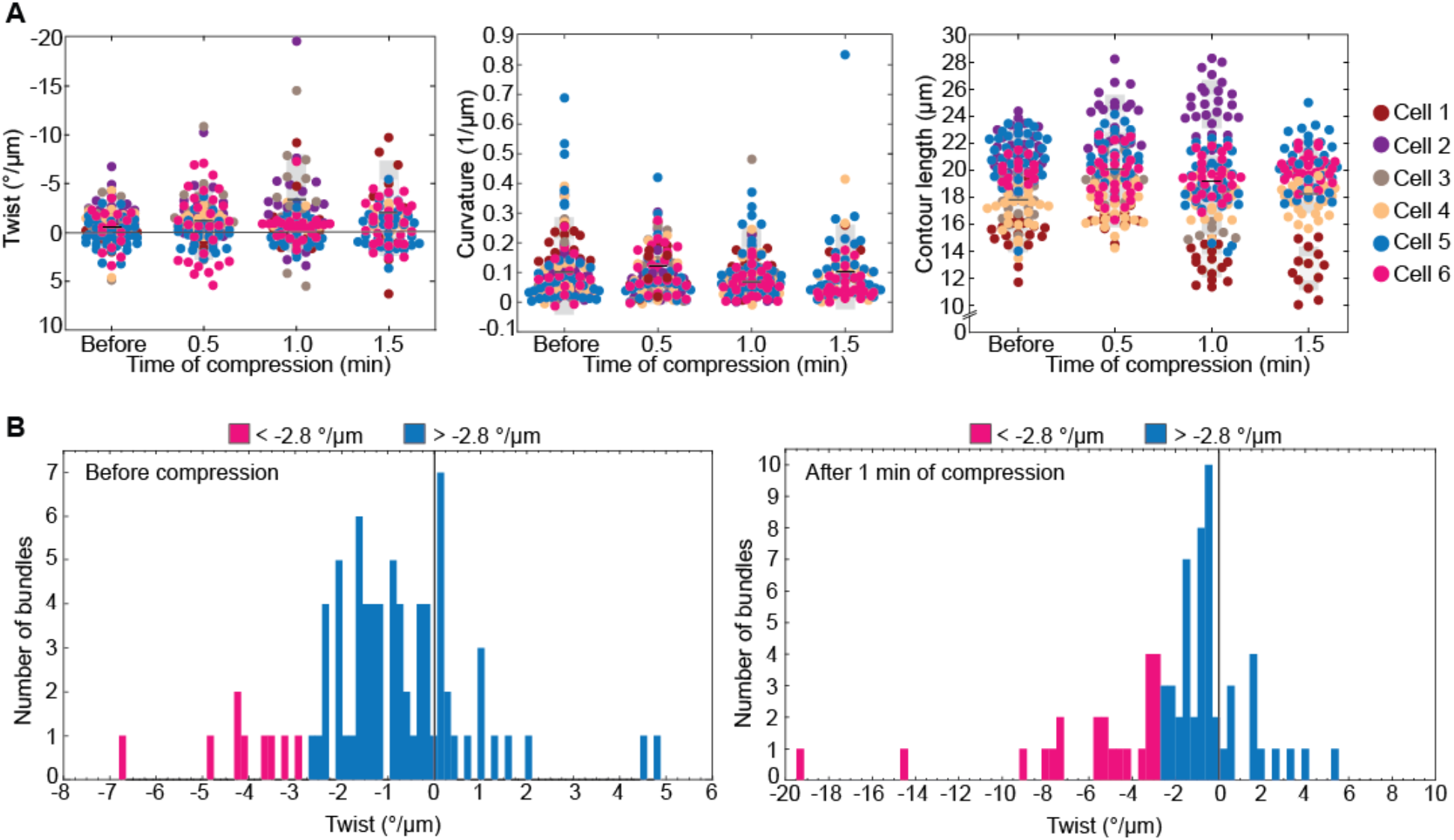
Twist, curvature and contour length of microtubule bundles in spindles compressed by an external force. **(A)** On the left, graph show the change of the twist before and up to 1.5 min after the compression. In the middle, graph shows the change of the curvature before and up to 1.5 min after the compression. On the right, graph shows the change of length of the bundle contours before and up to 1.5 min after the compression. Each color represents one cell, as described in the legend; dots represent bundles; the black line shows the mean; the light and dark grey areas mark 95% confidence interval on the mean and standard deviation, respectively. Experiments were performed on the HeLa-Kyoto BAC cells stably expressing PRC1-GFP. **(B)** Histograms of twist values before (left) and 1 minute after the compression (right). Colors magenta and blue represent bundles with twist values lower than −2.8 °/μm and above −2.8 °/μm (one standard deviation away from the mean twist before compression), respectively. Note that the distribution shifted towards more negative values upon compression. The twist was smaller than −2.8 °/μm (corresponding to strong left-handed twist) for 9 out of 80 bundles (11.3% ± 3.5%) before compression, whereas after compression this was the case for 21 out of 73 bundles (28.8% ± 5.3%).

**Figure S3.**
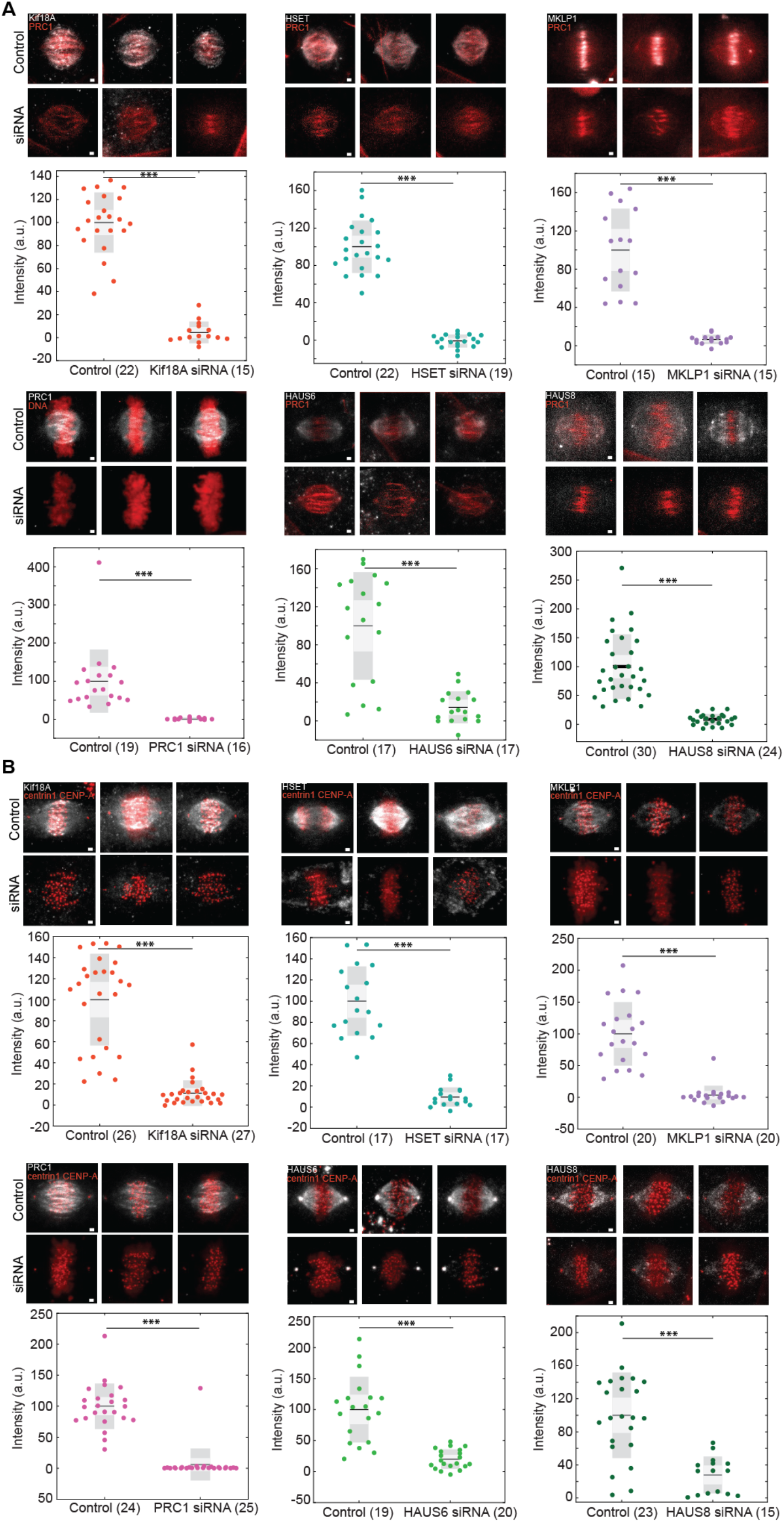
Immunofluorescence images of spindles in RPE1 and HeLa cells after protein perturbations. **(A)** Immunofluorescence of HeLa cells’ spindles after perturbation of spindle-associated proteins. Examples of three spindles for every perturbation of spindle-associated proteins and their controls: Kif18A depletion, MKLP1 depletion, HSET depletion, PRC1 depletion, HAUS6 and HAUS8 depletion, in that order. Microtubule bundles are shown in grey (proteins of interest) and DNA (dyed with DAPI in nontransfected HeLa) or PRC1 (in HeLa-Kyoto BAC cells stably expressing PRC1-GFP) in red. Images are shown in maximum z-projections. Graphs show intensities of protein of interest in control cells and cells treated with siRNA. ***p<0.001 (Student’s t-test). Experiments were performed on the non-transfected HeLa cell line (for the depletion of PRC1 and its control) and HeLa-Kyoto BAC cells stably expressing PRC1-GFP (for the rest of the treatments). **(B)** Immunofluorescence of RPE1 cells’ spindles after perturbation of spindle-associated proteins. Examples of three spindles for every perturbation of spindle-associated proteins and their controls: Kif18A depletion, HSET depletion, MKLP1 depletion, PRC1 depletion, HAUS6 and HAUS8 depletion, in that order. Microtubule bundles are shown in grey (proteins of interest) and kinetochores/centrosomes in red. Images are shown in maximum z-projections. Graphs show intensities of protein of interest in control cells and cells treated with siRNA. ***p<0.001 (Student’s t-test). Experiments were performed on the hTERT-RPE1 cells, permanently transfected and stabilized using CENP-A-GFP and centrin1-GFP.

**Figure S4.**
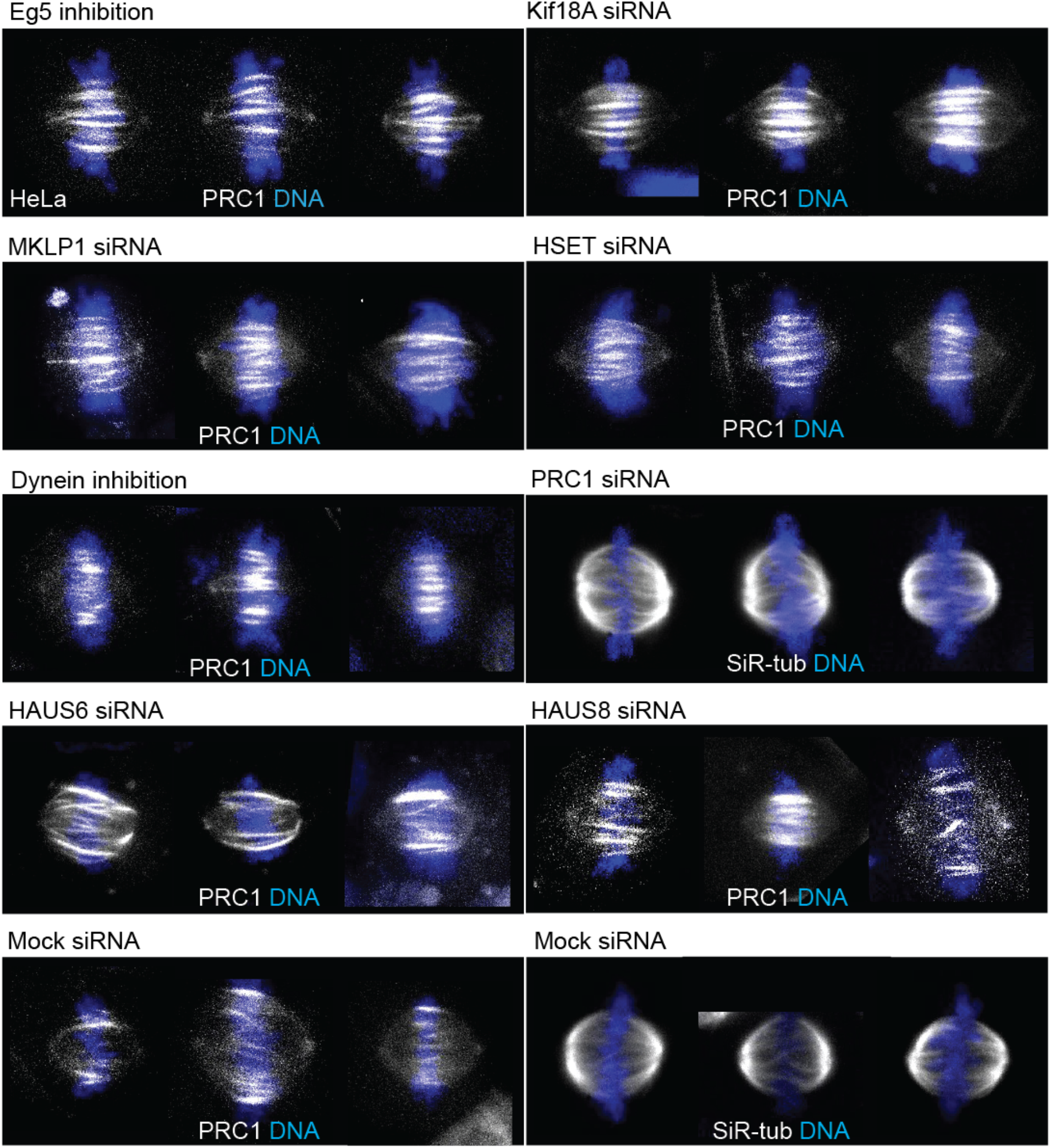
Spindles of HeLa cells after perturbation of spindle-associated proteins. Examples of three spindles for every perturbation of spindle-associated proteins: Eg5 inhibition, Kif18A depletion, MKLP1 depletion, HSET depletion, dynein inhibition, PRC1 depletion, HAUS6 and HAUS8 depletion and mock controls. Data was used in the graphs in Figs. 3B and 4B. Microtubule bundles are shown in grey (SiR-tubulin in non-transfected HeLa and PRC1-GFP in HeLa-Kyoto BAC cell line) and DNA in blue (SiR-DNA dye in HeLa-Kyoto BAC cell line and NucBlue dye in non-transfected HeLa). Images are shown in maximum z-projections. Experiments were performed on the non-tranfected HeLa cell line and HeLa-Kyoto BAC cells stably expressing PRC1-GFP.

**Figure S5.**
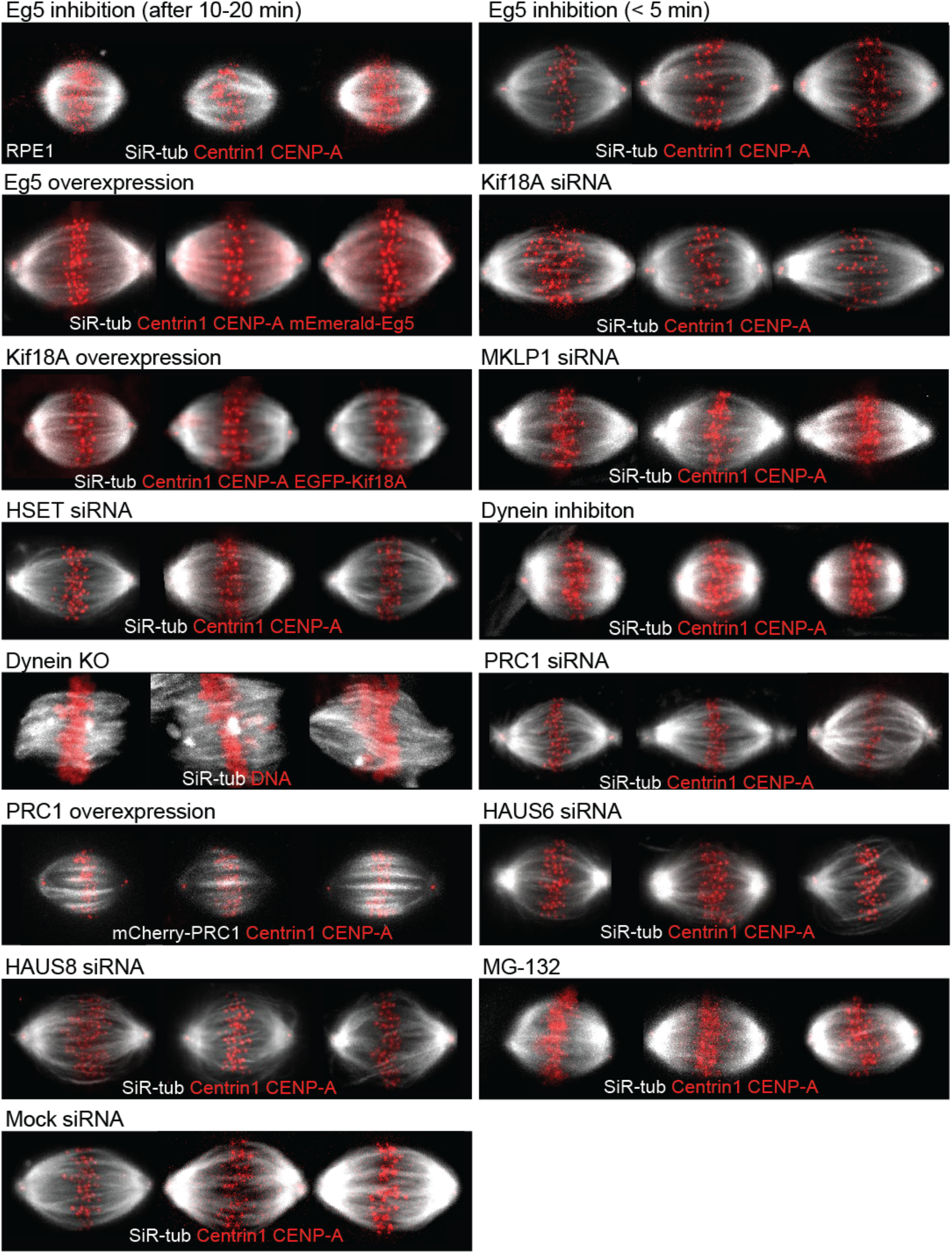
Spindles of RPE1 cells after perturbation of spindle-associated proteins. Examples of three spindles for every perturbation of spindle-associated proteins: Eg5 inhibition (after 10-20 min of STLC treatment), Eg5 inhibition (STLC treatment shorter than 5 min), Eg5 overexpression, Kif18A depletion, Kif18A overexpression, MKLP1 depletion, HSET depletion, dynein inhibition, dynein KO, PRC1 depletion, PRC1 overexpression, HAUS6 and HAUS8 depletion, MG-132 treatment, and mock control. Data was used in the graphs in Fig. 3C and 4C. Microtubule bundles are shown in grey (SiR-tubulin and, for PRC1 overexpression, mCherry-PRC1) and kinetochores/centrosomes, DNA (NucBlue dye), Eg5 and KiF18A in red. Images are shown in maximum z-projections. Experiments were performed on hTERT-RPE1 cells, permanently transfected and stabilized using CENP-A-GFP and centrin1-GFP and RPE1 inducible CRISPR/Cas9 DYNC1H1 knockout cells.

**Figure S6.**
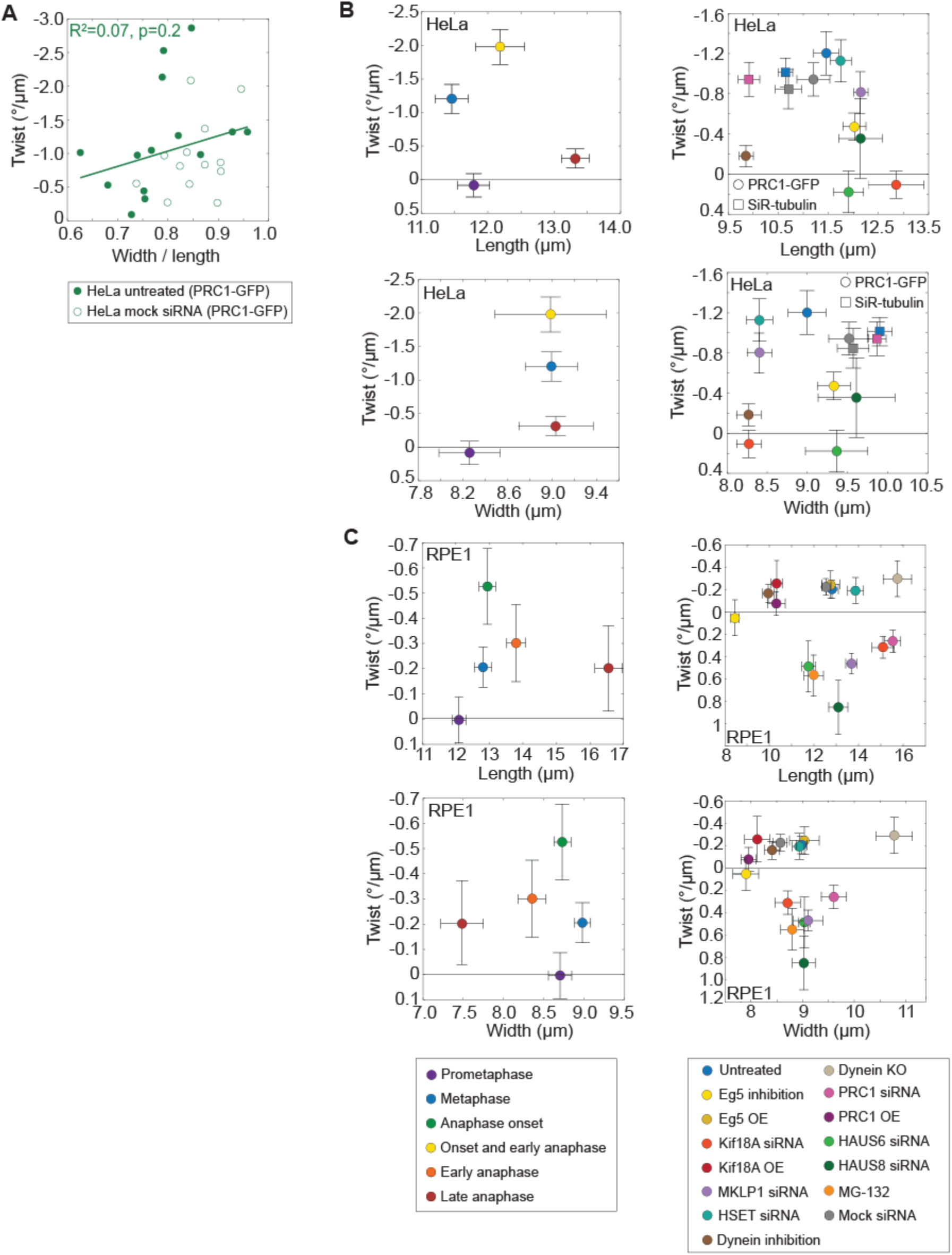
Correlation between spindle length or width and spindle twist in different phases of mitosis and during different protein perturbations in HeLa and RPE1 cells. **(A)** Round spindles have stronger twist than elongated spindles. Graph shows the correlation between width/length ratio and twist in HeLa cells. Filled circles represent untreated cells while empty circles represent mock siRNA controls. Lines show linear fit (untreated cells together with mock siRNA controls); equation y=-2.28x+0.79; goodness of fit shown in the graph. Same data was used in Figs. 1D, 3B and 4B. Experiments were performed on the HeLa-Kyoto BAC cells stably expressing PRC1-GFP. **(B)** Graphs showing how the spindle twist and length (top) or width (bottom) in HeLa cells vary depending on the different phases of mitosis (left) or perturbations of spindle-associated proteins (right). Each color represents different phase of mitosis or protein perturbation, as described in the legend at the bottom; error bars represent SEM. Same data was used in graphs in Fig. 1D, 3B and 4B. Experiments were performed on the HeLa-Kyoto BAC cells stably expressing PRC1-GFP (dots) and non-tranfected HeLa cell line for the depletion of PRC1 and its controls (rectangles). **(C)** Graphs showing how the spindle twist and length (top) or width (bottom) in RPE1 cells vary depending on the different phases of mitosis (left) or perturbations of spindle-associated proteins (right). Each color represents different phase of mitosis or protein perturbation, as described in the legend at the bottom; error bars represent SEM. Same data was used in graphs in Fig. 1E, 3C and 4C. Experiments were performed on hTERT-RPE1 cells, permanently transfected and stabilized using CENP-A-GFP and centrin1-GFP and RPE1 inducible CRISPR/Cas9 DYNC1H1 knockout cells.

### MOVIE CAPTIONS

**Movie S1:** Left-handed twist of the mitotic spindle in human cell. On the left, scheme of end-on point of view of the spindle; plane shows movement through z-planes. On the right, end-on view of fixed HeLa cell expressing PRC1-GFP (PRC1-GFP signal not shown); arrowheads point at the microtubule bundles that twist in a clockwise direction (indication of left-handed twist); blue asterisks represent spindle poles. Microtubule bundles are shown in grey; anti-α-tubulin in HeLa cell. Image of the spindle is shown in Fig. 1A.

**Movie S2:** Twist of prometaphase spindles. On the left, side-view (top) and end-on view (bottom) of prometaphase spindle in HeLa cell expressing PRC1-GFP; microtubule bundles are shown in grey (PRC1-GFP) and DNA in blue (SiR-DNA dye). On the right, side-view (top) and end-on view (bottom) of prometaphase spindle in RPE1 cells expressing CENP-A-GFP and centrin1-GFP; microtubule bundles are shown in grey (SiR-tubulin dye) and kinetochores/centrosomes (CENP-A-GFP/centrin-1-GFP) in red. Movies of spindles shown from the side view are played once while movies of spindles shown from the end-on view are repeated three times. Scale bar, 1 μm. Images of spindles are shown in Fig. 1B.

**Movie S3:** Twist of metaphase spindles. On the left, side-view (top) and end-on view (bottom) of metaphase spindle in HeLa cell expressing PRC1-GFP; microtubule bundles are shown in grey (PRC1-GFP) and DNA in blue (SiR-DNA dye). On the right, side-view (top) and end-on view (bottom) of metaphase spindle in RPE1 cells expressing CENP-A-GFP and centrin1-GFP; microtubule bundles are shown in grey (SiR-tubulin dye) and kinetochores/centrosomes (CENP-A-GFP/centrin-1-GFP) in red. Movies of spindles shown from the side view are played once while movies of spindles shown from the end-on view are repeated three times. Scale bar, 1 μm. Images of spindles are shown in Fig. 1B.

**Movie S4:** Twist of spindles at the beginning of anaphase. On the left, side-view (top) and end-on view (bottom) of the spindle at the beginning of anaphase in HeLa cell expressing PRC1-GFP; microtubule bundles are shown in grey (PRC1-GFP) and DNA in blue (SiR-DNA dye). On the right, side-view (top) and end-on view (bottom) of the spindle at the beginning of anaphase in RPE1 cells expressing CENP-A-GFP and centrin1-GFP; microtubule bundles are shown in grey (SiR-tubulin dye) and kinetochores/centrosomes (CENP-A-GFP/centrin-1-GFP) in red. Movies of spindles shown from the side view are played once while movies of spindles shown from the end-on view are repeated three times. Scale bar, 1 μm. Images of spindles are shown in Fig. 1B.

**Movie S5:** Twist of early anaphase spindles. On the left, side-view (top) and end-on view (bottom) of early anaphase spindle in HeLa cell expressing PRC1-GFP; microtubule bundles are shown in grey (PRC1-GFP) and DNA in blue (SiR-DNA dye). On the right, side-view (top) and end-on view (bottom) of early anaphase spindle in RPE1 cells expressing CENP-A-GFP and centrin1-GFP; microtubule bundles are shown in grey (SiR-tubulin dye) and kinetochores/centrosomes (CENP-A-GFP/centrin-1-GFP) in red. Movies of spindles shown from the side view are played once while movies of spindles shown from the end-on view are repeated three times. Scale bar, 1 μm. Images of spindles are shown in Fig. 1B.

**Movie S6:** Twist of late anaphase spindles. On the left, side-view (top) and end-on view (bottom) of late anaphase spindle in HeLa cell expressing PRC1-GFP; microtubule bundles are shown in grey (PRC1-GFP) and DNA in blue (SiR-DNA dye). On the right, side-view (top) and end-on view (bottom) of late anaphase spindle in RPE1 cells expressing CENP-A-GFP and centrin1-GFP; microtubule bundles are shown in grey (SiR-tubulin dye) and kinetochores/centrosomes (CENP-A-GFP/centrin-1-GFP) in red. Movies of spindles shown from the side view are played once while movies of spindles shown from the end-on view are repeated three times. Scale bar, 1 μm. Images of spindles are shown in Fig. 1B.

**Movie S7:** Twist before and after spindle compression. On the left, end-on view of the spindle in HeLa cell expressing PRC1-GFP before compression. On the right, end-on view of the spindle in HeLa cell expressing PRC1-GFP after 1 min of compression. Microtubule bundles are shown in grey (PRC1-GFP). Movies are played three times consecutively. Scale bar, 1 μm. Traces representing microtubule bundles are shown in Fig. 2B.

**Movie S8:** Twist of spindles in cells with depleted Kif18A. On the left, side-view (top) and end-on view (bottom) of the spindle in HeLa cell, expressing PRC1-GFP, after the depletion of Kif18A; microtubule bundles are shown in grey (PRC1-GFP) and DNA in blue (SiR-DNA dye). On the right, side-view (top) and end-on view (bottom) of the spindle in RPE1 cells, expressing CENP-A-GFP and centrin1-GFP, after the depletion of Kif18A; microtubule bundles are shown in grey (SiR-tubulin dye) and kinetochores/centrosomes (CENP-A-GFP/centrin-1-GFP) in red. Movies of spindles shown from the side view are played once while movies of spindles shown from the end-on view are repeated three times. Scale bar, 1 μm. Images of spindles are shown in Fig. 3A.

**Movie S9:** Twist of spindles in cells with depleted MKLP1. On the left, side-view (top) and end-on view (bottom) of the spindle in HeLa cell, expressing PRC1-GFP, after depletion of MKLP1; microtubule bundles are shown in grey (PRC1-GFP) and DNA in blue (SiR-DNA dye). On the right, side-view (top) and end-on view (bottom) of the spindle in RPE1 cells, expressing CENP-A-GFP and centrin1-GFP, after depletion of MKLP1; microtubule bundles are shown in grey (SiR-tubulin dye) and kinetochores/centrosomes (CENP-A-GFP/centrin-1-GFP) in red. Movies of spindles shown from the side view are played once while movies of spindles shown from the end-on view are repeated three times. Scale bar, 1 μm. Images of spindles are shown in Fig. 3A.

**Movie S10:** Twist of spindles in cells with perturbed dynein. On the left, side-view (top) and end-on view (bottom) of the spindle in HeLa cell, expressing PRC1-GFP, after the inhibition of dynein; microtubule bundles are shown in grey (PRC1-GFP) and DNA in blue (SiR-DNA dye). In the middle, side-view (top) and end-on view (bottom) of the spindle in RPE1 cells, expressing CENP-A-GFP and centrin1-GFP, after inhibition of dynein; microtubule bundles are shown in grey (SiR-tubulin dye) and kinetochores/centrosomes (CENP-A-GFP/centrin-1-GFP) in red. On the right, side-view (top) and end-on view (bottom) of the spindle in RPE1 inducible DYNC1H1 knockout cell, after the knockout of dynein heavy chain; microtubule bundles are shown in grey (SiR-tubulin dye) and DNA (NucBlue dye) in red. Movies of spindles shown from the side view are played once while movies of spindles shown from the end-on view are repeated three times. Scale bar, 1 μm. Images of spindles are shown in Fig. 4A.

**Movie S11:** Twist of spindles in cells with perturbed PRC1. On the left, side-view (top) and end-on view (bottom) of the spindle in non-tranfected HeLa cell after the depletion of PRC1; microtubule bundles are shown in grey (SiR-tubulin dye) and DNA in blue (NucBlue dye). In the middle, side-view (top) and endon view (bottom) of the spindle in RPE1 cells, expressing CENP-A-GFP and centrin1-GFP, after the depletion of PRC1; microtubule bundles are shown in grey (SiR-tubulin dye) and kinetochores/centrosomes (CENP-A-GFP/centrin-1-GFP) in red. On the right, side-view (top) and end-on view (bottom) of the spindle in RPE1 cells, expressing CENP-A-GFP and centrin1-GFP, after the overexpression of PRC1; microtubule bundles are shown in grey (PRC1-mCherry) and kinetochores/centrosomes (CENP-A-GFP/centrin-1-GFP) in red. Movies of spindles shown from the side view are played once while movies of spindles shown from the end-on view are repeated three times. Scale bar, 1 μm. Images of spindles are shown in Fig. 4A.

**Movie S12:** Twist of spindles in cells with depleted HAUS8. On the left, side-view (top) and end-on view (bottom) of the spindle with average twist value in HeLa cell, expressing PRC1-GFP, after the depletion of HAUS8; microtubule bundles are shown in grey (PRC1-GFP) and DNA in blue (SiR-DNA dye). In the middle, side-view (top) and end-on view (bottom) of the spindle with high right-handed twist in HeLa cell, expressing PRC1-GFP, after the depletion of HAUS8; microtubule bundles are shown in grey (PRC1-GFP) and DNA in blue (SiR-DNA dye). On the right, side-view (top) and end-on view (bottom) of the spindle in RPE1 cells, expressing CENP-A-GFP and centrin1-GFP, after the depletion of HAUS8; microtubule bundles are shown in grey (SiR-tubulin dye) and kinetochores/centrosomes (CENP-A-GFP/centrin-1-GFP) in red. Movies of spindles shown from the side view are played once while movies of spindles shown from the end-on view are repeated three times. Scale bar, 1 μm. Images of spindles are shown in Fig. 4A (except the spindle in HeLa cell with high right-handed twist).

## Notes

### Competing Interest Statement

The authors have declared no competing interest.

